# Astrocytes control quiescent NSC reactivation via GPCR signaling-mediated F-actin remodeling

**DOI:** 10.1101/2024.03.11.584337

**Authors:** Kun-Yang Lin, Mahekta R. Gujar, Jiaen Lin, Wei Yung Ding, Jiawen Huang, Yang Gao, Ye Sing Tan, Xiang Teng, Low Siok Lan Christine, Pakorn Kanchanawong, Yusuke Toyama, Hongyan Wang

## Abstract

The transitioning of neural stem cells (NSCs) between quiescent and proliferative states is fundamental for brain development and homeostasis. Defects in NSC reactivation are associated with neurodevelopmental disorders. *Drosophila* quiescent NSCs extend an actin-rich primary protrusion toward the neuropil. However, the function of the actin cytoskeleton during NSC reactivation is unknown. Here, we reveal the fine F-actin structures in the protrusions of quiescent NSCs by expansion and super-resolution microscopy. We show that F-actin polymerization promotes the nuclear translocation of Mrtf, a microcephaly-associated transcription factor, for NSC reactivation and brain development. F-actin polymerization is regulated by a signaling cascade composed of G-protein-coupled receptor (GPCR) Smog, G-protein αq subunit, Rho1 GTPase, and Diaphanous (Dia)/Formin during NSC reactivation. Further, astrocytes secrete a Smog ligand Fog to regulate Gαq-Rho1-Dia-mediated NSC reactivation. Together, we establish that the Smog-Gαq-Rho1 signaling axis derived from astrocytes, a NSC niche, regulates Dia-mediated F-actin dynamics in NSC reactivation.

## Introduction

Tissue development and homeostasis rely on the ability of stem cells to switch between quiescent and proliferative states. The majority of neural stem cells (NSCs) in the adult mammalian brain remain in a quiescent state, which upon physiological stimulation, are reactivated for adult neurogenesis and neural regeneration(*1*). Increasing evidence suggests that defects in NSC reactivation may be associated with aging-related cognitive decline(*2*) and neurodevelopmental disorders, such as. microcephaly(*3*). Deciphering the molecular mechanisms underlying NSC reactivation will provide important insights into brain development and regeneration and may facilitate the development of therapeutics for the treatment of brain ageing, injury, and neurological disorders.

NSCs in the *Drosophila* larval brain have emerged as an ideal *in vivo* model to investigate the molecular mechanisms underlying the transition between quiescent and proliferative states. *Drosophila* NSCs are programmed to enter the quiescent state at the end of embryogenesis(*4*), after which they undergo reactivation (undergo cell cycle re-entry and growth) within approximately 24 hours in the early larval stages upon feeding(*4*) (Fig. 1A). Dietary amino acids stimulate the secretion of insulin-like peptides in the brain-blood barrier (BBB) glial cells, an NSC niche in *Drosophila* larval brains; the insulin-like peptides activate the evolutionarily-conserved insulin/Insulin-like growth factor-1 (IGF-1) signaling(*5, 6*), which triggers NSC reactivation. Similar to *Drosophila* NSCs, mammalian NSCs are also activated by IGF-1 signaling, and mutations in the human IGF1 receptor (IGF1R) are linked to microcephaly(*7, 8*). In the mammalian brain, astrocytic glia produce Insulin-like growth factor 1 (IGF-1) for the proliferation of NSCs(*9*),(*10*). In addition, intrinsic factors required for NSC reactivation have also been identified, such as spindle matrix proteins(*11*), striatin-interacting phosphatase and kinase (STRIPAK) family proteins(*12*), Hsp83(*13*), CRL4^Mahj^ E3 ubiquitin ligase(*14*), Pr-set7(*15*), and microtubule-binding proteins Msps and Arf1(*16, 17*). On the contrary, the activation of the evolutionarily-conserved Hippo pathway keeps NSCs in quiescence(*18, 19*).

**Figure 1.**
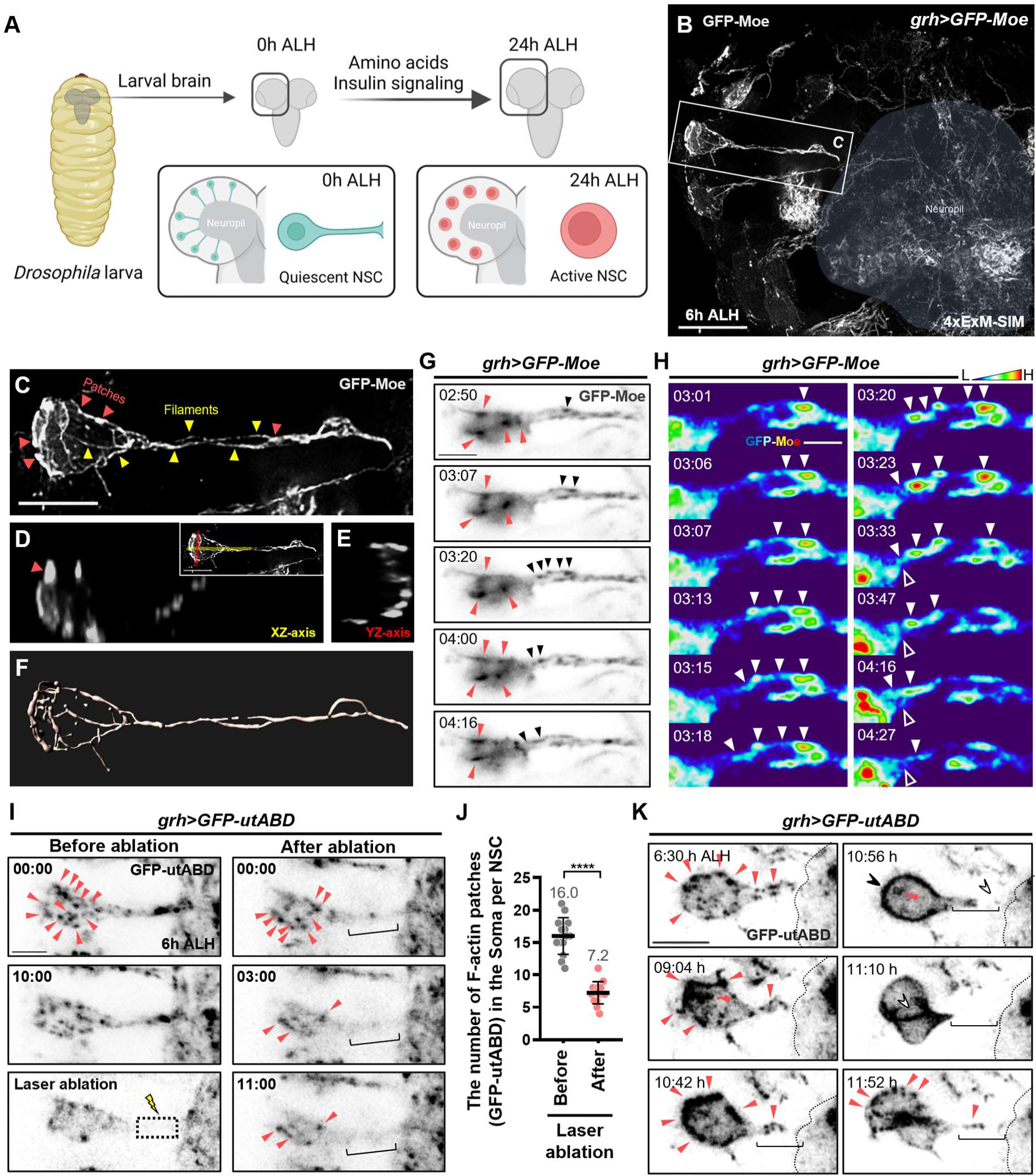
F-actin structures and dynamics during quiescent NSC reactivation. (**A**) Schematic diagram of quiescent NSC reactivation in the *Drosophila* larva brain. (**B**) Super-resolution imaging of F-actin structure achieved by ExM-SIM microscopy in *Drosophila* larval brain at 6h ALH. GFP-Moe (white) marks F-actin. Blue patches, neuropil region. (**C**) High magnification of quiescent NSC from **B**. Yellow arrowheads, F-filaments; red arrowheads, F-actin patches. (**D**)**-**(**E**) Orthogonal view (XZ-axis, E; YZ-axis, E’) of quiescent NSC in **C**. Insert of **D** shows sections used in **D** (XZ-axis in yellow) and **E** (YZ-axis in red). (**F**) 3D-reconstruction of D. (**G**) Live imaging stills of F-actin dynamics (GFP-Moe, black) in the quiescent NSC at 6h ALH. Time, mm:ss. F-actin patches, red arrowheads; black arrowheads, retrograde flow of F-actin patches in the protrusion. **(H**) Stills of live imaging of F-actin dynamics (GFP-Moe) in G in thermal theme. H for high and L for low in the thermal scale; white arrowheads, F-actin patches. Open arrowhead, PIS region. (**I**) Time-lapse images of GFP-utABD (black) driven by *grh-GAL4* were taken before and after laser ablation. Dash square, ablated area; red arrowheads, soma F-actin patches; brackets, the position of primary protrusion of quiescent NSCs. (**J**) Quantification graph of soma F-actin patches of quiescent NSCs before and after laser ablation. Before laser ablation: 16.0±2.8, n=14; after laser ablation: 7.2±1.7, n=14. Student *t*-test is used for statistics. ****P< 0.0001. The means of analyzed phenotypes were shown above each column. (**K**) Live imaging of GFP-utABD (black, F-actin) driven by *grh-GAL4*. Time, hh:mm. Red arrowheads, F-actin patches; black arrowhead, apical F-actin; white arrowhead, absence of F-actin. Bracket, primary protrusion. Dash lines indicate the boundary of neuropil. Scale bar, 10μm in B and K; 5μm in C, G and I; 2μm in H.

One hallmark of quiescent NSCs (qNSCs) in *Drosophila* is the presence of a primary cellular protrusion that extends from the cell body towards the neuropil. Recently, our laboratory demonstrated that this protrusion is enriched with the actin cytoskeleton(*11*). Although actin cytoskeleton is critical for the asymmetric division and cytokinesis of neural progenitors/stem cells(*20, 21*), actin dynamics in quiescent NSCs and their potential role in regulating NSC quiescence or reactivation have not been established. Diaphanous/Formin family proteins are key regulators of actin dynamics that accelerate F-actin nucleation and assembly(*22*). At the barbed end of the F-actin filaments, actin-polymerization factor Formins recruit and nucleate monomeric actin (also known as globular-actin or G-actin) for filament elongation(*22*). Interestingly, variants of Diaphanous-related formin 1 (DIAPH1) have been identified in human microcephaly patients(*23, 24*). Understanding whether and how Formin-mediated actin dynamics regulate NSC reactivation will provide new insights into developing therapeutic targets for neurodevelopmental disorders.

The heterotrimeric G protein complex is composed of three subunits, the α(Gα), β(Gβ) and γ(Gγ) subunits. Upon binding of ligand to the G-protein-coupled receptor (GPCR), Gα dissociates from the Gβγ subunits and gets activated. A dysfunction in GPCR signaling is associated with brain aging and neurodegenerative diseases(*25, 26*). Variants of GPCRs have also been identified in neurodevelopmental disorders including microcephaly(*27–29*). In human HEK293T cells, G protein alpha q subunit (Gαq) interacts with and regulates the activity of the small GTPase RhoA(*30*), which in turn, activates Formins to transduce extracellular stimuli into the assembly and organization of the actin cytoskeleton(*31*). Moreover, gain-of-function variants of GNAQ/Gαq have been identified in Sturge-Weber syndrome with neurological deficits, e.g., macrocephaly and seizures. Given the major roles of G protein signaling in fundamental cellular processes, the GPCR family has become a major drug target for treatments of various human diseases; 34% of FDA-approved drugs target GPCRs(*32*). Therefore, understanding how GPCR signaling controls neural stem cell reactivation may provide a potential strategy for the treatment of neurodevelopmental disorders.

Here, we unveiled the fine structure of the actin cytoskeleton and the retrograde flow of F-actin patches in quiescent NSCs. We showed that GPCR Smog-Gαq signaling regulates qNSC reactivation through Rho1-Diaphanous (Dia)-mediated actin dynamics. Moreover, the microcephaly-like phenotype of *dia* mutants could be suppressed by overexpression of the transcription factor Mrtf that is controlled by actin dynamics. We further identified astrocyte-like glia as a new NSC niche to produce Fog, a ligand of GPCR Smog, for qNSC reactivation. Our study demonstrates a new paradigm of NSC reactivation through astrocyte-mediated activation of GPCR signaling and regulation of actin dynamics.

## Results

### The dynamics of F-actin filaments and patches in quiescent neural stem cells

Previously, we showed that filamentous actin (F-actin) is enriched in the primary protrusion of quiescent NSCs (qNSCs)(*11*), however, the dynamics, structure, and function of the actin cytoskeleton in quiescent NSCs are unknown. To examine the structure of F-actin in qNSCs, we expressed the F-actin binding protein GFP-Moesin (GFP-Moe) using an NSC-specific GAL4 driver, *grainy head(grh)-GAL4*, in the NSCs of the *Drosophila* larva brain. As it is challenging to observe the fine structures (4-5 μm in diameter) of F-actin in qNSCs, we imaged GFP-Moe using structured illumination microscopy (SIM)(*33*). We further enhanced imaging resolution by using expansion microscopy (ExM)(*34*) for isotropic expansion of the fluorescent signal along with the expansion of sample structures (Fig. 1B). Compared with conventional confocal microscopy(*11*), ExM-SIM imaging greatly improved the resolution of F-actin structures in quiescent NSCs (Fig. 1C). Notably, F-actin was observed as long, twisted filaments in the primary protrusion (yellow arrowheads) and shorter filaments in the soma of qNSCs (Fig. 1C, yellow arrowheads). In addition, the soma also contained F-actin patches along the filaments (Fig. 1C, red arrowheads). Our orthogonal view (Fig. 1, D and E) and 3D reconstruction images (Fig. 1F and movie S1) pointed out a predominantly cortical localization of F-actin in the soma of quiescent NSCs. Next, we used live-cell imaging to monitor F-actin dynamics, marked by GFP-Moe, in qNSCs. Remarkably, at 6h ALH, F-actin patches in the soma underwent robust remodeling (Fig. 1G, red arrowheads and movie S2), as F-actin patches in the primary protrusion appeared to move in a retrograde flow towards the soma (Fig. 1H, thermal video stills, white arrowheads, stills 03:01 to 03:18 and 03:47 to 04:27; movie S2).

To understand whether this dynamics in the protrusion is important for F-actin polymerization in the soma of qNSCs, we examined F-actin dynamics labelled by GFP::utrophin-actin binding domain (GFP::utABD), followed by the severing of the protrusion using pico laser-induced ablation (see Methods for laser setting)(*17*). As expected, GFP::utABD was not recovered at 10 mins post-injury (Fig.1I), as opposed to its rapid recovery in the fluorescence recovery after photobleaching (FRAP) experiment (Fig S1, A and B; 50% recovery at 100 seconds after photobleaching). Remarkably, F-actin patches in the soma diminished dramatically after injury to the protrusion (Fig. 1, I and J; movie S3), suggesting that F-actin patches might move from the protrusion back to the soma in qNSCs.

### Dynamics of the actin cytoskeleton during quiescent NSC reactivation

Actin dynamics during the asymmetric division of proliferative NSCs has been well characterized(*20, 21, 35, 36*). However, F-actin dynamics during the switch between quiescent and proliferative states is unknown. As GFP-Moe did not survive beyond 24h ALH, likely due to toxicity of GFP-Moe overexpression, GFP::utABD that survived to adult stages was used to mark F-actin dynamics during quiescent NSC reactivation. Up to 10h ALH, F-actin patches in the primary protrusion moved retrogradely toward the soma (Fig. 1K, 6:30-10:42 hh:mm ALH; movie S4), similar to the retrograde flow seen using GFP-Moe-marked F-actin in quiescent NSCs. Strikingly, GFP::utABD signal diminished from the distal end of the protrusion (10:56 hh:mm ALH, white arrowhead) and subsequently disappeared from the entire protrusion (10:42-10:56 hh:mm ALH), which was associated with a dramatic reduction in the number of F-actin patches in the soma of NSCs (red arrows) and an enrichment of F-actin at the apical region of the NSCs (Fig. 1K, black arrows; 10:56 hh:mm ALH; movie S4). As the NSC re-entered into the cell cycle to generate two daughter cells (Fig. 1K, 11:10 hh:mm ALH; movie S4), F-actin accumulated in the cleavage furrow, as expected, during cytokinesis (Fig. 1K, 11:10, white arrow). Following the division, F-actin patches reappeared in the soma and the protrusion, with the latter attached to the presumptive basal daughter cell-ganglion mother cell (Fig. 1K, 11:52; movie S4). While the protrusion in qNSCs persisted throughout the first cell division, it underwent a thinning process(*17, 37*), possibly due to loss of F-actin patches during the reactivation.

Taken together, our ExM-SIM imaging has clearly demonstrated the presence of fine F-actin structures in quiescent NSCs. Our live imaging analysis suggested that F-actin patches can move from the protrusion to the soma in quiescent NSCs.

### Depletion of *diaphanous* causes microcephaly-like phenotype and defects in quiescent NSC reactivation

To identify actin regulators that are involved in quiescent NSC reactivation, we performed a small-scale RNAi screen (unpublished) and found that Diaphanous (Dia), the sole Formin in *Drosophila*, is required for NSC reactivation. At 24 h ALH, the majority of quiescent NSCs in control larval brains re-entered into cell cycle and incorporated EdU, with only 17.9% of NSCs remaining quiescent and negative for EdU (Fig. 2, A and B). In contrast, the percentage of quiescent NSCs that were EdU-negative notably increased to 42.7%-69.5% in two loss-of-function alleles, *dia^1^* and *dia^5^,* and two hemizygous *dia* mutants, *dia^1/Df(2L)ED1317^* and *dia*^5/^*^Df(2L)ED1317^* (Fig. 2, A and B). In addition, the percentage of quiescent NSCs carrying primary protrusions, the hallmark of quiescent NSCs, increased significantly from 5.8% in the control to 15.1% and 16.3% in the *dia^1^* and *dia^5^* mutant brains, respectively (Fig. 2, C and D). Dia protein levels were diminished in the cytosol of *dia^5^* mutant NSCs and *dia*-knocked down NSCs, while Dia was sequestered in the nucleus of *dia^1^* mutant NSCs, resulting in a mild reduction of Dia in the cytoplasm (Fig. S2, A to C). Variants of *Diaphanous-Related Formin 1(DIAPH1)*, the orthologue of *Drosophila dia* in humans, have been identified in microcephaly patients(*23, 24, 38*). Consistent with this, the volumes of *dia^1^* and *dia^5^* brain lobes were dramatically reduced to 28% and 37.5%, respectively, compared with the control (100%) at 96h ALH (Fig. 2, E and F), suggesting that *Drosophila dia* mutants also display a microcephaly-like phenotype. *dia* RNA interference (RNAi) in NSCs under the control of *grh-GAL4* also resulted in defects in NSC reactivation (Fig. 2, G to I), suggesting that Dia is required intrinsically in the NSCs for their reactivation. Likewise, knocking down *dia* via another NSC driver, *inscuteable-GAL4* driver (*insc-GAL4*), resulted in NSC reactivation defect (Fig. S4, A and B). In contrast, *dia* knockdown in glial cells or fat body did not cause any reactivation defect (Fig. S5). Our results suggest that the actin polymerization factor Dia is a novel intrinsic regulator of quiescent NSC reactivation and brain development.

**Figure 2.**
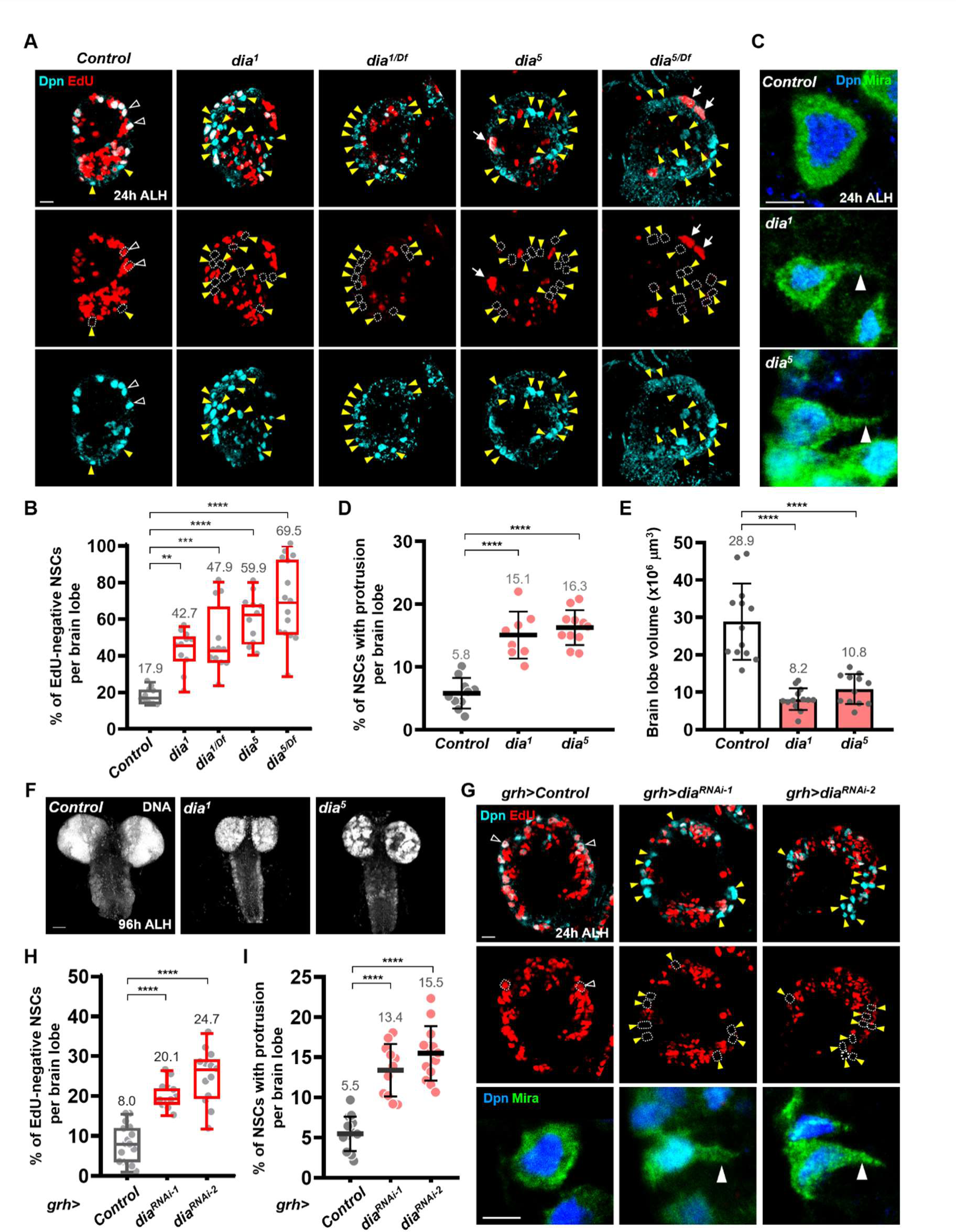
The actin polymerization factor Diaphanous/Formin is required for quiescent NSC reactivation and brain development. (**A**) Larval NSCs were labelled with EdU and Dpn at 24h ALH. Yellow arrowheads and dash circles point to EdU-negative NSCs. White arrowheads, EdU-positive NSCs. White arrows, NSCs with cytokinesis defects. (**B**) The quantification graph of EdU-negative NSCs in (A). *Control (yw)*: 17.9±4.4, n=10; *dia^1^*: 42.7±10.9, n=11; *dia^1/Df^*: 47.9±18.0, n=12; *dia^5^*: 59.9±13.1, n=11; *dia^5/Df^*: 69.5±21.6, n=15. (**C**) NSCs were stained for Dpn and Mira. White arrowheads point to primary protrusion of quiescent NSCs. (**D**) Quantification graph of the percentage of NSCs carrying primary protrusion. *Control (yw)*: 5.8±2.4, n=10; *dia^1^*: 15.1±3.7, n=8; *dia^5^*: 16.3±2.8, n=11. (**E**) The quantification graph of brain volume. *Control (yw)*: 28.9±10.2, n=13; *dia^1^*: 8.2±2.9, n=13; *dia^5^*: 10.8 ± 4.0, n=12. (**F**) the size of larval brains (DNA, gray). (**G**) Top and middle rows, proliferating NSCs (Dpn, cyan; EdU, red) in control (*β-gal^RNAi^*) or *dia*-KD larval brains (driven by *grh*-GAL4). Yellow arrowheads, EdU-negative NSCs; white open arrowheads, EdU^+^ proliferative NSCs. Bottom row, NSCs labelled with Dpn/Mira. White solid arrowheads, primary protrusion of quiescent NSCs. (**H**) The quantification graph of EdU-negative NSCs. *Control* (*β-gal^RNAi^)*: 8.0±4.7, n=16; *dia^RNAi-1^*: 20.1±3.3, n=12; *dia^RNAi-2^*: 24.7±6.7, n=13. **I,** Quantification graph of the percentage of NSCs retaining primary protrusion. *Control* (*β-gal^RNAi^)*: 5.5±2.1, n=14; *dia^RNAi-1^*: 13.4±3.3, n=11; *dia^RNAi-2^*: 15.5±3.4, n = 13. One-Way ANOVA is used for statistics. ****P<0.0001; ***P< 0.001; **P< 0.01. The means of analyzed phenotypes were shown above each column. Scale bar, 50μm in F; 10μm in A, G (top row); 5μm in C and G (bottom row).

### Heterotrimeric G protein Gαq is required for NSC reactivation

It is well known that G-protein-coupled receptor (GPCR)/G-protein signaling regulates actin dynamics to control many cellular processes, such as morphogenesis, cell motility and axonal development(*39*). To assess the role of GPCR/G-protein signaling in qNSC reactivation, we performed an RNAi screen on various heterotrimeric G protein subunits (unpublished) and identified the G alpha q subunit (Gαq) as a new regulator of NSC reactivation, as *Gαq* knockdown results in defects in qNSC reactivation (Fig. S3, A to C). Importantly, qNSCs in larval brains carrying loss-of-function alleles of *Gαq* (*Gαq^221C^*) and a hemizygous *Gαq^221C/Df(2R)Gαq1.3^* mutant had a delayed reactivation (36.9% EdU-negative NSCs in *Gαq^221C^* and 36.8% EdU-negative NSCs in *Gαq^221C/ Df(2R)Gαq1.3^*) compared to control (14.9% EdU-negative NSCs) at 24 h ALH (Fig. 3, A and C; Fig. S3, D and E). In addition, more NSCs retained primary protrusions in *Gαq^221C^*-mutant larval brains than in the control (Fig. 3, A and D). However, knocking down *Gαq* in glial cells or fat body did not cause any reactivation defect (Fig. S5). These observations suggest that Gαq is required intrinsically for qNSC reactivation.

**Figure 3.**
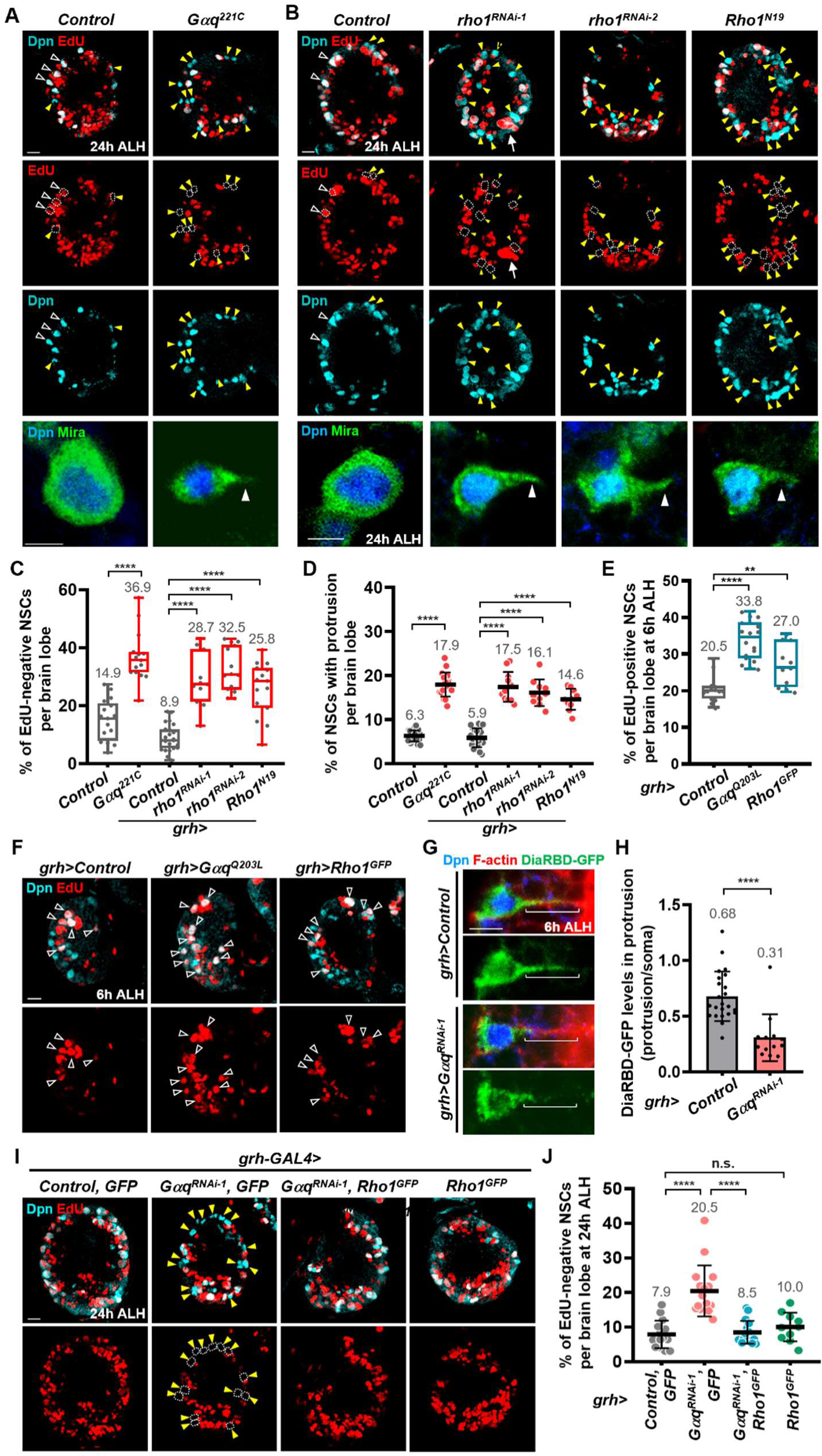
Gαq-Rho1 signaling promotes quiescent NSC reactivation. (**A**) **and (B)** Larval NSCs were labelled with EdU and Dpn at 24h ALH. Yellow arrowheads and dash circles, EdU-negative NSCs; White arrowheads, EdU-positive NSCs. Bottom row, NSCs were labelled with Dpn and Mira. White arrowheads, protrusion of NSCs. White arrows, NSCs with cytokinesis defects. (**C**) Quantification graph of EdU-negative quiescent NSCs in A-B. *Control (yw)*: 14.9±7.1, n=17; *Gαq^221C^*: 36.9±8.8, n=15; *Control* (*β-gal^RNAi^)*: 8.9±4.6, n=23; *rho1^RNAi-1^*: 28.7±9.3, n=11; *rho1^RNAi-2^*: 32.5±7.7, n=14*; Rho1^N19^*: 25.8±9.2, n=15. (**D**) Quantification graph of NSCs retaining protrusion in A-B. *Control (yw)*: 6.3±1.3, n=21; *Gαq^221C^*: 17.9±2.7, n=18; *Control* (*β-gal^RNAi^)*: 5.9±2.1, n=20; *rho1^RNAi-1^*: 17.5 ± 3.4, n=11; *rho1^RNAi-2^*: 16.1±3.0, n=10*; Rho1^N19^*: 14.6±2.4, n=10. (**E**) Quantification graph of EdU-positive NSCs at 6h ALH. *Control (β-galRNAi)*: 20.5±3.6, n=14; *Gαq^Q203L^*: 33.8±5.3, n=18; *Rho1^GFP^*: 27.0±5.8, n=11. (**F**) Proliferating NSCs (EdU, red; Dpn, cyan) in control (*β-gal^RNAi^*), *Gαq^Q203L^*, *GFP-Rho1* larval brains driven by *grh-GAL4* at 6h ALH. White arrowheads, EdU^+^ NSCs. (**G**) Quiescent NSCs (Dpn, cyan) in control (*β-gal^RNAi^*), *Gαq*-KD larval brains under *grh-GAL4* at 6h ALH. F-actin (Rhodamine-Phalloidin) marks protrusions. DiaRBD-GFP (green) marks active Rho1. (**H**) Quantification graph of DiaRBD-GFP levels in the protrusion in (G). *Control* (*β-gal^RNAi^)*: 0.68±0.22, n=23; *Gαq^RNAi-1^*: 0.31±0.21, n=13. (**I**) Proliferating NSCs (EdU, red; Dpn, cyan) in larval brains at 24h ALH. UAS-GFP, a negative control for suppression effect. Yellow arrowheads and dash circles, EdU-negative NSCs. (**J**) Quantification graph of EdU-negative NSCs in (I). *Control (β-gal^RNAi^),GFP*: 7.9±4.0, n=15; *Gαq^RNAi-1^, GFP*: 20.5±7.4, n=16; *Gαq^RNAi-1^, Rho1^GFP^*: 8.5±3.3, n=16; *Rho1^GFP^*: 10.0±4.1, n=10. One-Way ANOVA (**C, D, E, J**) and unpaired student t test (**H**) are used for statistics. ****P<0.0001; ***P<0.001; **P<0.01. The means of analyzed phenotypes were showed above each column. Scale bar, 10μm in A, B (top row), F and I; 5μm in A, B (bottom row) and G (bottom row).

Although Dia is known to be required for cytokinesis (*40*), no cytokinesis defects were found in *dia^1^* mutant larval brains at 24h ALH (n=11), and *dia* knockdown only exhibited a very mild cytokinesis defect in NSCs (5%, n=10) at 24h ALH. Even in *dia^5^*, the null mutant, only a weak cytokinesis defect (Fig. 2A, white arrows; 11%, n=10) in NSCs was observed at 24h ALH. In addition, *rho1* knockdown resulted in a negligible cytokinesis phenotype (3%, n=11), while Gαq knockdown had no obvious effect on NSC cytokinesis (n=15). Therefore, the cytokinesis defect does not significantly contribute to the reactivation phenotypes observed in this study. In addition, *Gαq* and *dia* knockdown using *grh-GAL4* did not obviously affect larval growth or the pupariation rate.

To investigate whether Gαq activation is sufficient to promote NSC reactivation upon fed and nutrition restriction, we overexpressed a constitutively active form of Gαq (Gαq^Q203L^) under the control of *grh-GAL4*. Remarkably, Gαq^Q203L^ overexpression triggered an increase in the number of NSCs re-entering the cell cycle (Fig. 3, E and F; control: 20.5% EdU-positive NSCs; Gαq^Q203L^: 33.8%) only on fed condition, but not on nutrition restriction condition (Fig. S6).

### Small GTPase Rho1 is required for NSC reactivation

Small GTPase Rho1/RhoA is a known activator of Dia/Formin(*41*) and a downstream effector of Gαq signaling that controls F-actin remodeling in fibroblast (NIH 3TC) and human embryonic kidney (HEK) 293 cells (*30, 42–44*). Therefore, we investigated the function of *Drosophila* Rho1 in qNSC reactivation. Since loss-of-function alleles of *Rho1* causes embryonic lethality, we analyzed the effect of Rho1 knockdown in the larval brain. In two independent *rho1* RNAi strains under the control of *grh*-Gal4, the percentage of NSCs without EdU incorporation was significantly increased from 8.9% in the control NSCs to 28.7% and 32.5 % in *rho1^RNAi-1^* (BDSC #9909)- and *rho1^RNAi-2^* (BDSC #9910)-expressing NSCs, respectively, at 24 h ALH (Fig. 3, B and C). Likewise, overexpression of a dominant-negative form of Rho1 (Rho1^N19^), driven by *grh-GAL4,* resulted in an increased percentage of EdU-negative NSCs (25.8%) at 24 h ALH (Fig. 3, B and C). In addition, more NSCs retained primary protrusions in *rho1^RNAi-1^*-, *rho1^RNAi-2^*- or *Rho1^N19^*-expressing NSCs (Fig. 3, B and D). These data established an intrinsic role for Rho1 in quiescent NSC reactivation. Similar to Gαq^Q203L^ overexpression, overexpression of GFP-Rho1 in NSCs resulted in a significant increase in the number of reactivated NSCs (Fig. 3, E and F) on fed condition. Our observations suggest that Rho1 is required for NSC cell cycle re-entry in the presence of nutrition.

### Gαq promotes quiescent NSC reactivation through Rho1 activation

To assess the cellular pattern of active Rho1 in quiescent NSCs, we took advantage of Dia-Rho1-binding domain (DiaRBD)-GFP, which binds specifically to active Rho1(*45*), and overexpressed it in the NSCs using *grh-GAL4* driver. DiaRBD-GFP was found in both the soma and primary protrusions of control qNSCs at 6h ALH (Fig. 3, G and H). Strikingly, Gαq knockdown resulted in a specific reduction in DiaRBD-GFP signal in the primary protrusions (Fig. 3, G and H), suggesting Gαq-mediated Rho1 activation in the primary protrusions of qNSCs. Next, we determined whether Gαq acts upstream of Rho1 during NSC reactivation. Indeed, GFP-Rho1 overexpression fully suppressed defects in cell cycle re-entry caused by Gαq depletion (Fig. 3, I and J). While there were 20.5% EdU-negative NSCs in *Gαq*-KD NSCs (with GFP-overexpression as a control) at 24 h ALH, GFP-Rho1 overexpression reduced the number of quiescent NSCs to 8.5% (n=16) upon *Gαq* depletion, which was indistinguishable from the control group (Fig. 3, H and I; 7.9%, n=15). Meanwhile, the overexpression of GFP-Rho1 alone did not affect cell cycle re-entry of qNSCs at 24 h ALH (Fig. 3, I and J). These results strongly suggest that Gαq promotes cell cycle re-entry of quiescent NSCs through Rho1 activation in the primary protrusion.

### Gαq-Rho1 signaling promotes NSC reactivation via Dia-mediated F-actin remodeling

Given that Gαq is required for Rho1 activation in the primary protrusions of quiescent NSCs (Fig. 3G), we determined whether Gαq is also required for Dia protein localization in these cells. Using anti-Dia antibodies, we detected Dia in the cytoplasm of soma and primary protrusions of quiescent NSCs (Fig. 4, A and B). Remarkably, knocking down *Gαq* using two independent RNAi lines dramatically reduced Dia levels in the primary protrusions (Fig. 4, A and B). Similarly, Dia localization in the primary protrusion was diminished upon *rho1* depletion in NSCs (Fig. 4, A and B).

**Figure 4.**
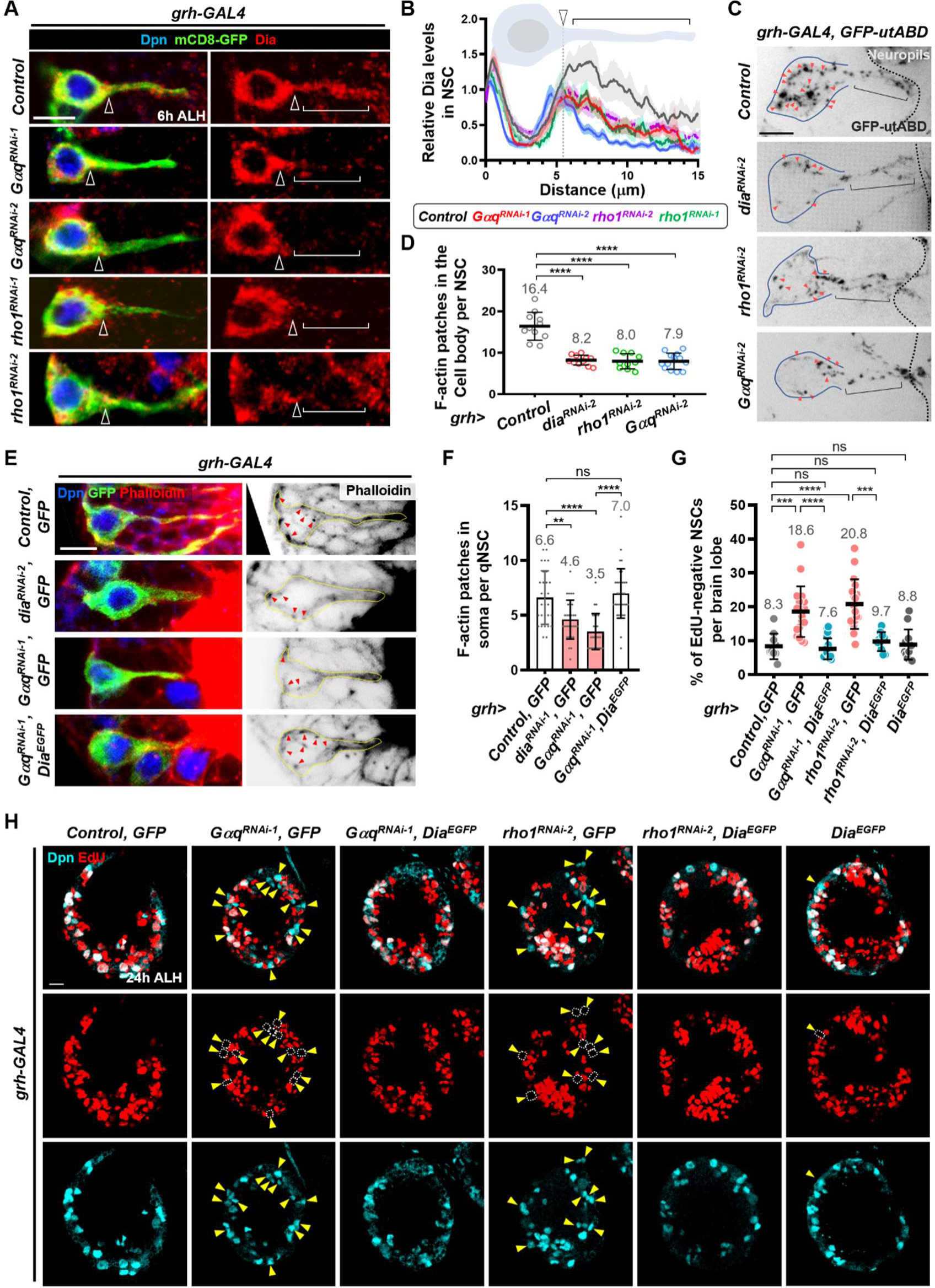
Gαq-Rho1-Dia signaling promotes NSC reactivation via actin cytoskeleton. (**A**) Dia protein (red) in quiescent NSCs (Dpn, blue; mCD8-GFP, green) in control (*β-gal^RNAi^*), *Gαq*-KD, *rho1*-KD larval brains under the control of *grh*-GAL4 driver at 6h ALH. Yellow arrowheads, neck region of quiescent NSC. Brackets, primary protrusion. (**B**) Quantification graph of Dia protein levels along the soma and protrusion of quiescent NSC in various RNAi transgenes driven by *grh-GAL4* driver. (**C**) F-actin patches (GFP-utABD, black) in control *(β-gal^RNAi^*), *Gαq*-KD, *rho1*-KD and *dia*-KD quiescent NSCs under the control of *grh-GAL4* driver at 6h ALH. Solid lines outline the soma of NSCs. Dash lines indicate the boundary of neuropil. Brackets, primary protrusion. (**D**) Quantification graph of F-actin patches in the soma of quiescent NSCs in various genotypes of C. *Control* (*β-gal^RNAi^)*: 16.4±3.4, n=11; *dia^RNAi-2^*: 8.2±1.2, n=11; *rho1^RNAi-2^*: 8.0±1.8, n=16; *Gαq^RNAi-2^*: 7.9±1.9, n=13. (**E**) F-actin (Rhodamine-Phalloidin, red and black) in quiescent NSCs (Dpn, blue; mCD8-GFP, green) under *grh-GAL4* driver at 6h ALH. Red arrowheads, F-actin patches. Yellow lines mark intact quiescent NSCs. (**F**) Quantification graph of F-actin patches in the soma of quiescent NSCs in various genotypes of G. *Control* (*β-gal^RNAi^),GFP*: 6.6±2.4, n=29; *dia^RNAi-1^, GFP*: 4.6±1.8, n=29; *Gαq^RNAi-1^, GFP*: 3.5±1.6, n=25; *Gαq^RNAi-1^, Dia^EGFP^*: 7.0±22, n=34. (**G**) Quantification graph of EdU-negative quiescent NSCs in various genotypes of F. *Control* (*β-gal^RNAi^),GFP*: 8.3±3.8, n=10; *Gαq^RNAi-1^, GFP*: 18.6±7.5, n=18; *Gαq^RNAi-1^, Dia^EGFP^*: 7.6±3.1, n=19; *rho1^RNAi-1^,GFP*: 20.8±7.3, n=18; *rho1^RNAi-1^, Dia^EGFP^*: 9.7±2.8, n=10; *Dia^EGFP^*: 8.8±4.5, n=13. (**H**) Proliferating NSCs (EdU, red; Dpn, cyan) in larval brains of various transgenes driven by *grh-GAL4* at 24h ALH in G. Yellow arrowheads and dash circles point to EdU-negative NSCs. One-Way ANOVA is used for statistics. ****P<0.0001; ***P<0.001; **P<0.01. The means of analyzed phenotypes were shown above each column. Scale bar, 10μm in H; 5μm in A, C and E.

Next, we assessed the effects of Gαq, Rho1 and Dia on F-actin remodeling in quiescent NSCs at 6h ALH by live-cell imaging of GFP::utABD. F-actin retrograde flow from the primary protrusion to the soma was significantly disrupted in the *dia*-, *rho1*- and *Gαq*-KD qNSCs (Fig. S7; movie S5). In addition, the depletion of these genes resulted in a dramatic decrease in the number of F-actin patches in the soma of qNSCs (Fig. 4, C and D; movie S6). These data suggest that the Gαq-Rho1-Dia signaling axis regulates the retrograde flow and polymerization of F-actin in the soma of quiescent NSCs.

Next, we performed epistatic analysis to determine whether Dia functions downstream of Gαq to mediate F-actin remodeling and/or NSC reactivation. Indeed, Dia-EGFP overexpression under *grh*-GAL4 completely suppressed the reduction in F-actin patches and defects in NSC reactivation in Gαq-depleted quiescent NSCs (Fig. 4, E to H). Dia-EGFP overexpression alone, in the absence of Gαq depletion, did not affect the reactivation (Fig. 4, G and H). Likewise, Dia-EGFP overexpression suppressed *rho1* KD-induced defects in NSC cell cycle re-entry (Fig. 4, G and H). Our results suggest that the Gαq-Rho1 signaling axis promotes NSC reactivation via Dia-mediated F-actin remodeling.

### F-actin polymerization is required for the reactivation of quiescent NSCs

Since Dia promotes F-actin polymerization and NSC reactivation, we determined whether F-actin polymerization is important for quiescent NSC reactivation. To address this, we examined the potential roles of two groups of F-actin polymerization regulators, the positive regulators Profilin (*chickadee* (*chic)* in *Drosophila*), *spire* (*spir)* and *singed* (*sn*) and the negative regulators Cofilin (*twinstar* (*tsr)* in *Drosophila*), *capping protein beta* (*cpb)* and *capulet (capt)*. A reduction in F-actin polymerization via knocking down positive regulators or overexpressing negative regulators results in defective NSC reactivation (Fig. S8A). In contrast, when F-actin polymerization was enhanced by the depletion of negative regulators or overexpression of positive regulators, quiescent NSCs reactivation was unaltered (Fig. S8B). Interestingly, knocking down Arp2, Arp3 or WASp, regulators for F-actin branching, in NSCs did not cause NSC reactivation defect (Fig. S8C), suggesting that branched F-actin is not important for NSC reactivation. These findings strongly indicate that F-actin polymerization is required for NSC reactivation.

### The Hippo-Wts-Yki signaling pathway is not a functional target of Dia in NSC reactivation

To understand how F-actin remodeling influence NSC reactivation, we assessed whether Dia could control NSC reactivation via regulating Yki activity in quiescent NSCs. Some earlier studies have suggested that Yorkie (Yki) activity, which is controlled by the evolutionarily-conserved Hippo signaling pathway and which can respond to biomechanical cues such as tension within the F-actin cytoskeleton in various cell types(*46, 47*), is necessary for quiescent NSC reactivation(*18, 19*). Interestingly, nuclear Yki levels were significantly reduced in *dia^1^-* and *dia^5^-*mutant qNSCs (Fig. S9, A and B). Similarly, RNAi induced-*dia* knock down reduced nuclear Yki levels in the NSCs (Fig. S9, C and D). However, neither overexpression of a constitutively-active form of Yki (Yki^S168A^-GFP) nor knocking down *Warts* (*Wts)* that phosphorylates and inactivates Yki could suppress NSC reactivation defects in the *dia*-depleted larval brains (Fig. S9, E and F). These observations suggest that the reduction of nuclear Yki might be a consequence, but not the cause of delayed reactivation upon *dia*-depletion and that the Hippo-Wts-Yki signaling pathway is unlikely to be a functional target of Dia during NSC reactivation.

### Transcription factors Mrtf and SRF are required for NSC reactivation

Formin-mediated F-actin assembly triggers the nuclear translocation of the conserved myocardin-related transcription factor (MRTF) and activates MRFT/serum response factor (SRF) signaling in mammalian smooth muscle cells and embryonic fibroblasts(*48–51*). *Drosophila* Mrtf is known to regulate tracheal branching and cell migration(*52, 53*), but its potential function during brain development is not established. Thus, we examined the function of Mrtf and Blistered (Bs, *Drosophila* homolog of SRF) in NSC quiescence exit. Remarkably, NSC reactivation was impaired in both *mrtf* and *bs* mutants; quiescent NSCs in larval brains carrying loss-of-function alleles of *Mrtf^Δ7^* and a hemizygous *Mrtf^Δ7/Df(3L)BSC412^* showed delayed reactivation (51.4% EdU-negative NSCs in *Mrtf^Δ7^* larvae and 45.1% EdU-negative NSCs in *Mrtf^Δ7/Df(3L)BSC412^* larvae) compared with control (17.9% EdU-negative NSCs) at 24 h ALH (Fig. 5, A and C). Similarly, *mrtf* knockdown in NSCs led to delayed reactivation, seen by an increase in the percentage of EdU-negative NSCs from 9.8% in the control to 20.2% and 29% in the *mrtf^RNAi-1^*- and *mrtf^RNAi-2^*-expressing NSCs, respectively, at 24 h ALH (Fig. 5, B and C). The findings suggested that *mrtf* is intrinsically required for NSC reactivation. In addition, larval brains carrying loss-of-function alleles of *bs^Δ0326^* and a hemizygous *bs^03267/Df(2R)Exel6082^* showed delayed NSC reactivation (29.6% EdU-negative NSCs in *bs^Δ0326^* and 38.1% EdU-negative NSCs in *bs^03267/Df(2R)Exel6082^*larvae) compared with control NSCs (17.9% EdU-negative NSCs) at 24 h ALH (Fig. S10, A and B). These data suggest that SRF-Mrtf signaling promotes quiescent NSC reactivation.

**Figure 5.**
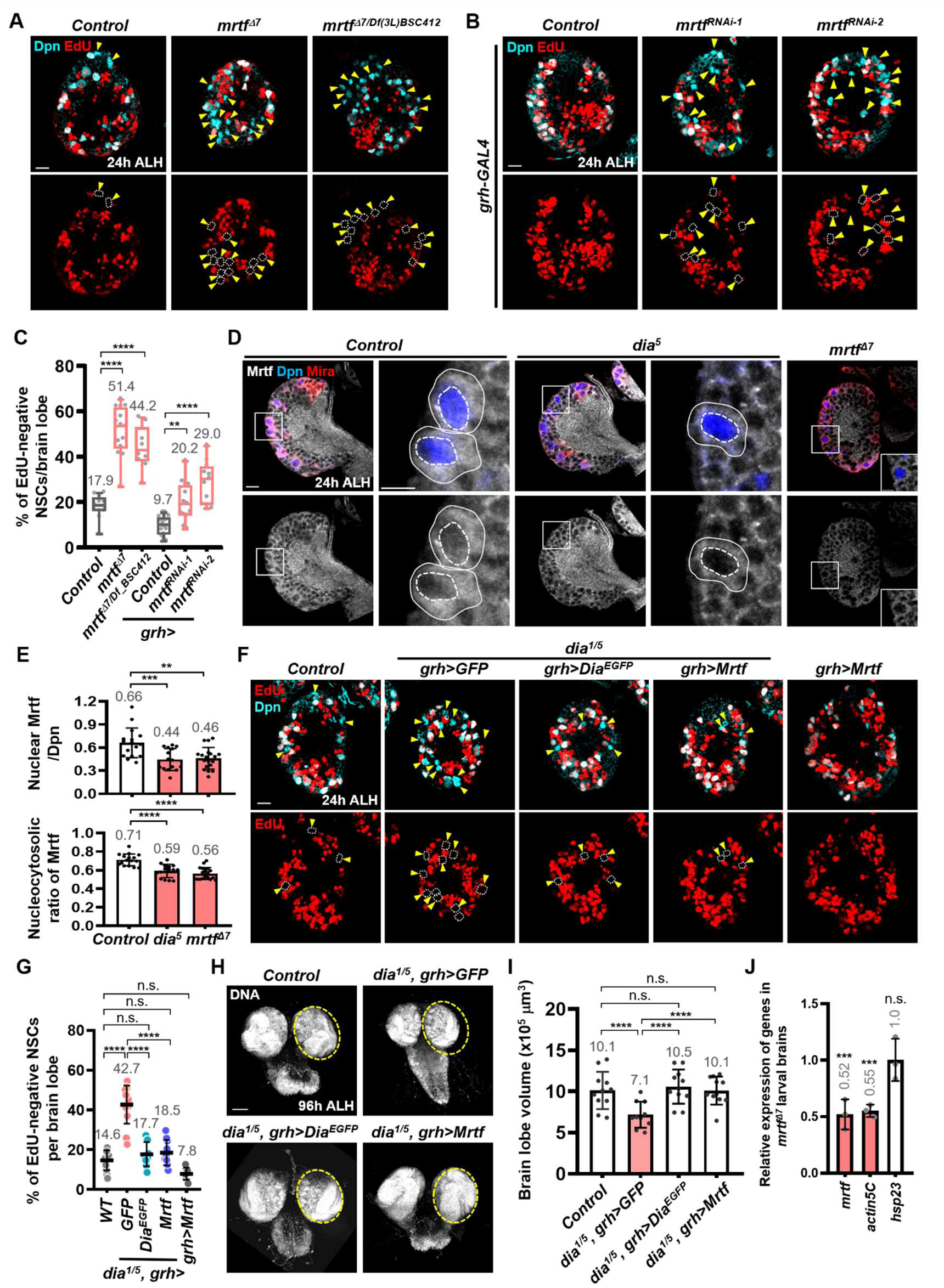
Diaphanous promotes quiescent NSC reactivation and brain development via transcription factor Mrtf. (**A**) and (**B**) Larval NSCs were labelled with EdU and Dpn at 24h ALH. Yellow arrowheads and dash circles, EdU-negative NSCs. (**C**) Quantification graph of EdU-negative NSCs in (A-B). *Control(yw)*: 17.9±5.2, n=11; *mrtf^D7^*: 51.4±10.7, n=14; *mrtf^D7/Df(3L)BSC412^*: 44.2±9.2, n=10. *Control* (*β-gal^RNAi^)*: 9.7±4.1, n = 15; *mrtf^RNAi-1^*: 20.2±8.7, n=13; *mrtf^RNAi-2^*: 29.0±9.0, n=11. (**D**) NSCs were labelled with Mrtf, Dpn and Mira. White squares, the region for high magnification. Solid lines, NSCs. Dash circles, nucleus. (**E**) Quantification graph of nuclear Mrtf expression (top) and ratio of nuclear Mrtf to cytoplasmic Mrtf (bottom). Top graph: *Control (yw)*: 0.66±0.2, n=16; *dia^5^*: 0.44±0.14, n=15; *mrtf^D7^*: 0.46±0.14, n=16. Bottom graph: *Control (yw)*: 0.71±0.06, n=16; *dia^5^*: 0.59±0.07, n=15; *mrtf^D7^*: 0.56±0.06, n=19. (**F**) Larval NSCs were labelled with EdU and Dpn at 24h ALH. Yellow arrowheads and dash circles, EdU-negative NSCs. (**G**) Quantification graph of EdU-negative NSCs in E. *Control (yw)*: 14.6±5.0, n=11; *dia^1/5^, grh>GFP*: 42.7±9.6, n=13; *dia^1/5^, grh>Dia^EGFP^*: 17.7±6.1, n=10; *dia^1/5^, grh>Mrtf*: 18.5±6.4, n=10; *grh>Mrtf*: 7.8 3.0, n=10. (**H**) The size of larval brains (DNA, gray) at 96h ALH. Dash circles, single brain lobe. (**I**) Quantification graph of brain size from *Control (yw)*: 10.1±2.3, n=11; *dia^1/5^, grh>GFP*: 7.1±1.6, n=10; *dia^1/5^, grh>Dia^EGFP^*: 10.6±2.1, n=11; *dia^1/5^, grh>Mrtf*: 10.1±1.7, n=11. (**J**) Quantification graph of RT-qPCR analysis in 24 h ALH brains from control (*yw*) and *mrtf^Δ7^*. After normalization against *yw* control (with SD): *mrtf*, 0.52±0.08-fold; *actin5C*, 0.55 ± 0.03-fold; *Hsp23*, 1.00 ± 0.11-fold. One-Way ANOVA is used for statistics. ***P<0.001; ns, no significance. The means of analyzed phenotypes were shown above each column. P<0.0001; ***P<0.001; **P<0.01. The means of analyzed phenotypes were shown above each column. Scale bar, 50μm in H; 10μm in A, B, D and F; 5μm in D for images with high magnification.

### Nuclear Mrtf localization correlates with the active state of NSCs

To assess the localization pattern of Mrtf in NSCs, we expressed GFP-tagged Mrtf, Mrtf-3xGFP (driven by ubiquitous *tubulin(tub)* promoter)—a functional form that can fully rescue the ovarian defect seen in the loss-of-function allele of *mrtf^Δ7^* (*54*). Notably, nuclear Mrtf-GFP intensity was higher in both mushroom body NSCs at 0h ALH and central brain NSCs at 24h ALH when compared with non-mushroom body NSCs at 0h ALH (Fig. S10, C to I), suggesting that nuclear Mrtf correlates with the active status of NSCs.

To examine the localization of endogenous Mrtf in NSCs, we generated polyclonal anti-Mrtf antibodies against C-terminal *Drosophila* Mrtf (1119–1418 amino acids encoded in exon 5). Endogenous Mrtf was localized in both the cytoplasm and nucleus in control NSCs (Fig. 5, D and E), similar to GFP-tagged Mrtf (Fig. S10, C to H). The nuclear intensity as well as the nucleocytoplasmic ratio of Mrtf were decreased in *Mrtf^Δ7^* mutant NSCs at 24h ALH (Fig. 5, D and E). In addition, total Mrtf protein was reduced in *Mrtf^Δ7^* and *Mrtf^KO^* mutant brains, in which exon 1 and exons 4-5 were deleted, respectively (Fig. S10, J and K), suggesting the specificity of the antibody.

### The Gαq-Rho1-Dia pathway promotes NSC reactivation through the transcription factor Mrtf

We wondered whether Dia would be important for the nuclear localization of Mrtf in NSCs via F-actin polymerization. Indeed, the nuclear intensity and nucleocytoplasmic ratio of Mrtf were significantly decreased in *dia^5^-*mutant NSCs (Fig. 5, D and E), as well as in *Gαq^221C^* mutant-, *rho1*-KD-, and *Rho1^N19^*-OE NSCs at 24h ALH (Fig. S11, A to D).

Next, we assessed whether Mrtf overexpression could suppress the *dia-*mutant phenotypes. Remarkably, Mrtf overexpression completely suppressed defects in EdU incorporation in the transheterozygous *dia* mutant (*dia^1/5^*) (Fig. 5, F and G), while overexpression of Mrtf alone did not affect NSC reactivation (Fig. 5, F and G). Likewise, Mrtf overexpression significantly suppressed defects in EdU incorporation in *Gαq*-KD and *rho1*-KD NSCs (Fig. S11, E and F). These results indicate that Mrtf acts downstream of the Gαq-Rho1-Dia pathway to promote quiescent NSC reactivation.

Similar to DIAPH1, variants in MRTF have been identified in human patients with microcephaly(*23, 24, 55, 56*). We found that overexpression of Mrtf in the NSCs reversed the microcephaly-like phenotype seen in the transheterozygous *dia^1/5^* mutant to the control brain volume (Fig. 5, H and I), further supporting that Mrtf functions downstream of Dia-mediated F-actin remodeling in the NSCs during brain development.

Since actin is a known transcriptional target of Mrtf in *Drosophila* ovary and breast cancer cells(*54*), we tested whether Mrtf could regulate *actin* in the larval brain. Interestingly, *actin5C* expression was significantly reduced in *mrtf^Δ7^* mutant brains (Fig. 5J), suggesting a positive feedback regulation between Mrtf and Actin that leads to enhanced *actin* expression and Mrtf nuclear localization during quiescent NSC reactivation.

### GPCR Smog promotes quiescent NSC reactivation via the Gαq-Dia pathway

To identify the GPCR that activates Gαq in the NSCs, we examined the expression of a total of 123 GPCRs from the published single-cell RNA seq database of the *Drosophila* larval brain(*57*). From this dataset, we identified 36 GPCRs that were expressed in NSCs (Fig. S12, A to C). Next, we performed a small-scale RNAi screen, and five out of 36 GPCRs showed potential defects in NSC reactivation. Among these five candidates, we found that a GPCR named Smog, a protein interacting with Gαq in the mouse brain(*58*), is required for quiescent NSC reactivation (Fig. 6, A and B). At 24 h ALH, two independent *smog* RNAi lines displayed defects in NSC reactivation with a significant increase in EdU-negative NSCs (20.5% in *smog^RNAi-1^* (BDSC #43135) and 25.1% in *smog^RNAi-2^* (BDSC #51705)) compared to control NSCs (7.9%) (Fig. 6, A and B). Importantly, quiescent NSCs in larval brains carrying a loss-of-function allele *smog^knock-out(KO)^* and a hemizygous mutant *smog^KO/Df(2L)Exel9062^* showed delayed NSC reactivation (41.6% EdU-negative NSCs in *smog^KO^* larvae and 33.9% EdU-negative NSCs in *smog^KO/Df(2L)Exel9062^* larvae compared to 16.7% EdU-negative NSCs in the control larvae) at 24 h ALH (Fig. 6, C and D). *smog* expression indicated by mCD8-GFP (under the control of *smog*-GAL4) was notably higher in quiescent NSCs than in active NSCs, supporting a role for Smog in the reactivation (Fig. S12, D and E). We examined the subcellular localization of Smog in quiescent NSCs using GFP-tagged Smog driven by *spaghetti squash*(*sqh)* promoter(*59*) and found it to be enriched in the primary protrusion of these cells (Fig. 6, E and F). Moreover, knocking down *smog* in quiescent NSCs in both *smog^RNAi-1^* and *smog^RNAi-2^* significantly reduced Dia protein levels in the primary protrusion (Fig. 6, G and H), similar to *Gαq* or *Rho1* depletion (Fig. 4, A and B).

**Figure 6.**
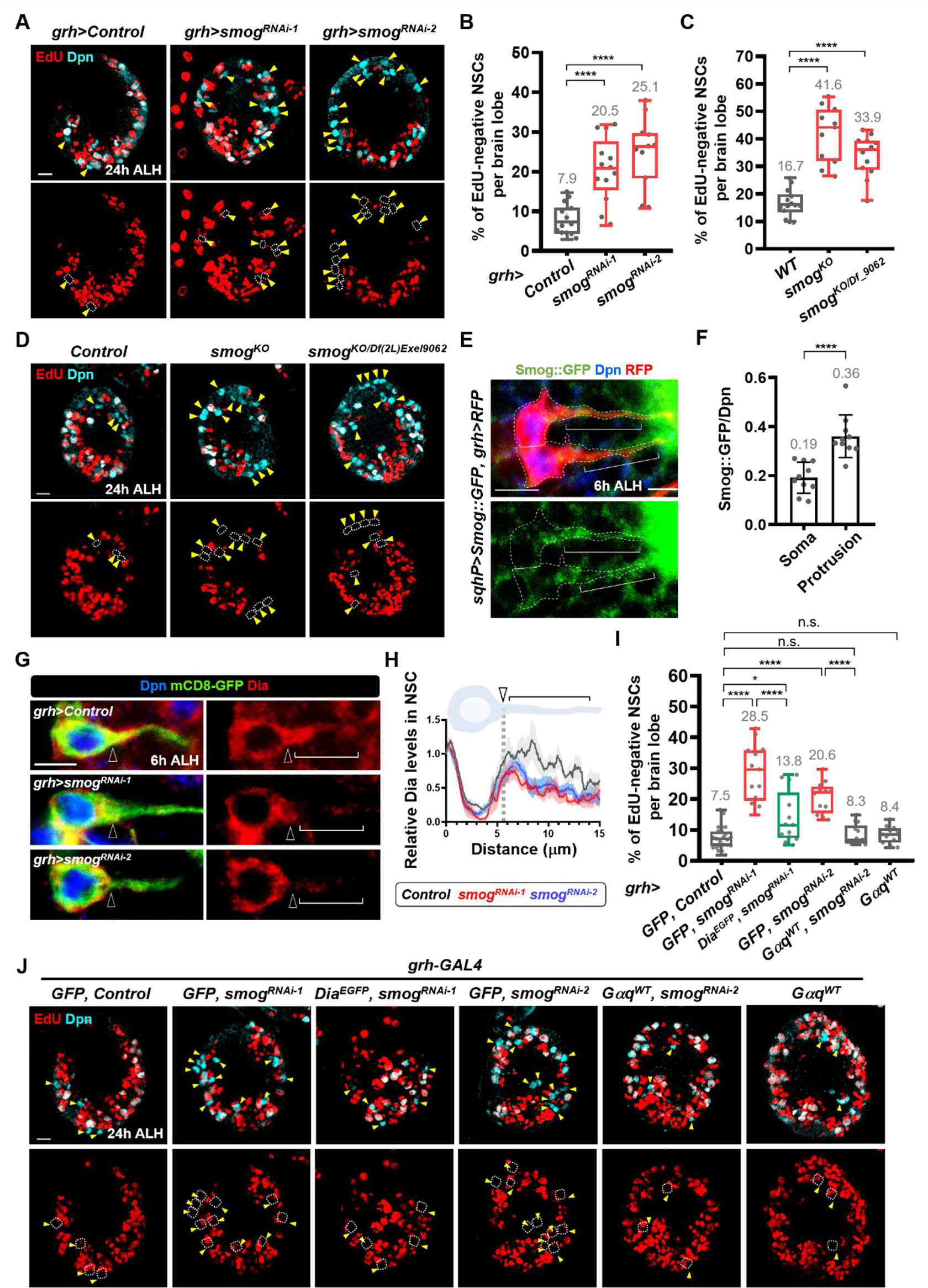
GPCR Smog promotes quiescent NSC reactivation via Gαq-Dia signaling. (**A**) Proliferating NSCs (EdU, red; Dpn, cyan) in control (*β-gal^RNAi^*) and *smog*-KD larval brains under the control of *grh-GAL4* driver at 24h ALH. Yellow arrowheads and dash circles point to EdU-negative NSCs. (**B**) Quantification graph of EdU-negative NSCs of A at 24h ALH. *Control* (*β-gal^RNAi^)*: 7.9±3.8, n=14; *smog^RNAi-1^*: 20.5±8.1, n=13; *smog^RNAi-2^*: 25.1±8.7, n=11. (**C**) Quantification graph of EdU-negative NSCs in control (*yw*) and *smog* mutant flies in D. *Control (yw)*: 16.7±4.9, n=14; *smog^KO^*: 41.6±10.0, n=11; *smog^KO/Df(2L)Exel9062^*: 33.9±7.6, n=12. (**D**) Proliferating NSCs (EdU, red; Dpn, cyan) in larval brains at 24h ALH. (**E**) Smog::GFP localization (green) in the quiescent NSCs (Dpn, blue; RFP; red) in the larval brains at 6h ALH. Dash lines, intact qNSCs. Brackets, primary protrusions. (**F**) Quantification graph of Smog::GFP levels in the soma and protrusion of quiescent NSCs. Soma: 0.19±0.06, n=10; protrusion: 0.36±0.09, n=10. (**G**) Dia protein (red) in control (*β-gal^RNAi^*) and *smog*-KD quiescent NSCs (Dpn, blue, mCD8-GFP, green) under the control of *grh-GAL4* driver at 6h ALH. (**H**) Quantification graph of Dia protein levels along the soma and protrusion in control (*β-gal^RNAi^*) and *smog*-KD quiescent NSCs at 6h ALH in G. (**I**) Quantification graph of EdU-negative NSCs in J. *Control* (*β-gal^RNAi^),GFP*: 7.5±3.7, n=23; *smog^RNAi-1^, GFP*: 28.5±9.2, n=13; *smog^RNAi-1^, Dia^EGFP^*: 13.8±8.3, n=13; *smog^RNAi-2^, GFP*: 20.6±5.1, n=12; *smog^RNAi-2^, Gαq^WT^*: 8.3±3.3, n=13 *Gαq^WT^*: 8.4±2.8, n=11. (**J**) Proliferating NSCs (EdU, red; Dpn, cyan) in larval brains at 24h ALH. Yellow arrowheads and dash circles, EdU-negative NSCs. One-Way ANOVA (**B, C, L**) and unpaired student t test (**F,H**) were used for statistics. ****P < 0.0001; ***P < 0.001; no significance. The means of analyzed phenotypes were shown above each column. Scale bar, 10μm in A, D, and J; 5μm in E and G.

To investigate whether Gαq and Dia act downstream of GPCR Smog to promote NSC reactivation, we overexpressed Dia^EGFP^ or wild type Gαq (Gαq^WT^), in the *smog*-KD NSCs. Remarkably, overexpression of Dia^EGFP^ or Gαq^WT^ in the *smog*-KD NSCs significantly suppressed the defect of EdU incorporation in NSCs (Fig. 6, I and J). Overexpressing Gαq^WT^ alone did not affect the reactivation of quiescent NSCs (Figure Fig. 6, I and J). These data suggest that GPCR Smog promotes quiescent NSC reactivation via the Gαq-Dia axis.

#### Astrocytes secrete Folded gastrulation (Fog) to promote quiescent NSC reactivation

Folded gastrulation (Fog) is a known ligand for the GPCR Smog in *Drosophila* mesoderm, salivary gland, and S2 cells(*59, 60*). Fog-Smog signaling controls Rho1 activity for epithelial tube formation during salivary gland invagination in the fly(*60*). However, the role of Fog during quiescent NSC reactivation is unknown. We sought to identify the cell type in the larval brain that secretes the Fog protein. We took advantage of a dataset of single cell RNA-sequencing (scRNA-seq) published by C. B. Avalos, et al(*57*) and analyzed *fog* expression in different cell types of the larval brain. Unexpectedly, *fog* was primarily expressed in the glial cells (Fig. 7A), but not in neural stem cells or neurons. Intriguingly, only a small proportion (∼20%) of glial cells had *fog* expression (Fig. 7A). This prompted us to pinpoint the subtype of glial cells that expresses *fog*. There are four subtypes of glial cells in the *Drosophila* central brain, including surface glia (perineurial glia and subperineurial glia), cortex glia, ensheathing glia, and astrocytes(*61*). Our analysis on the same scRNA-seq dataset revealed that *fog* is predominantly expressed in the astrocytes, and to much lesser levels in the ensheathing glia, but not in the surface glia or cortex glia (Fig. 7B). These analyses pointed out the astrocytes as the main glial cells secreting the Fog protein in the *Drosophila* larval brain.

**Figure 7.**
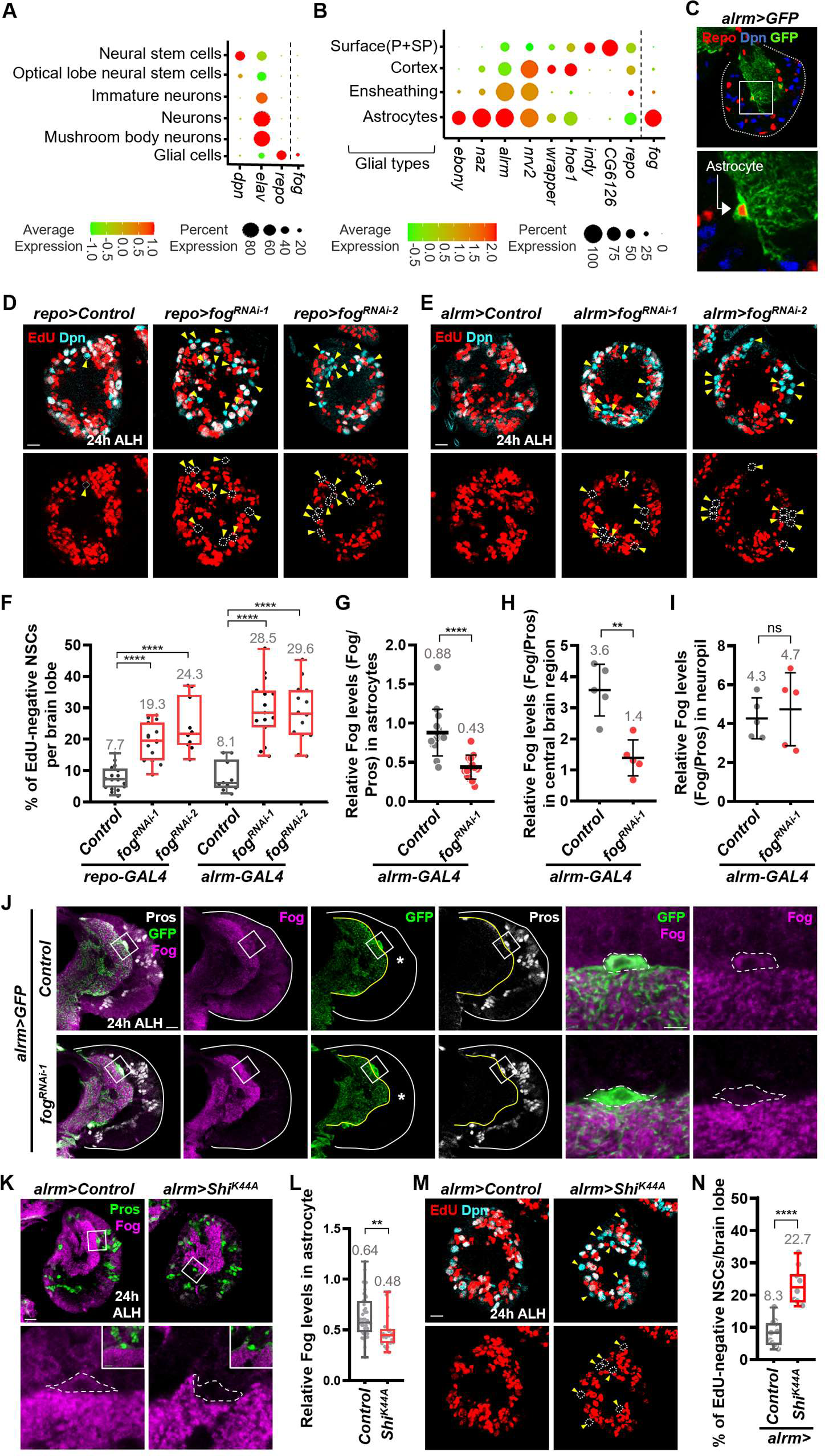
Astrocytes, a new NSC niche, secrete Fog via dynamin to promote quiescent NSC reactivation. (**A**) **and** (**B**) *fog* mRNA expression in larval brains from the dataset of single-cell RNA seq (refer to Materials and Methods). (**C**) Astrocyte (Repo and *alrm>GFP*) in larval brain at 24h ALH. (**D**) and (**E**) Larval NSCs (EdU and Dpn) at 24h ALH. Yellow arrowheads and dash circles, EdU-negative NSCs (**F**) Quantification graph of EdU-negative NSCs in C and D. *repo*>*control*: 7.7± 3.8, n=17; *repo*>*fog^RNAi-1^*: 19.3±5.8, n=15; *repo*>*fog^RNAi-2^*: 24.3±8.4, n=10; *alrm*>*control*: 8.1 ± 4.8, n=10; *alrm*>*fog^RNAi-1^*: 28.5±9.1, n=14; *alrm*>*fog^RNAi-2^*: 29.6±9.6, n=15. (**G**) Quantification graph of Fog levels in astrocytes: control, 0.88±0.29, n=15; *alrm>fog^RNAi-1^*, 0.43±0.15, n=14. (**H**) Fog levels in central brain region: control, 3.57±0.83, n=5; *alrm>fog^RNAi-1^*, 1.39±0.58, n=5. (**I**) Fog levels in neuropil region: control, 4.27±1.05, n=5; *alrm>fog^RNAi-1^*, 4.73±1.87, n=5. (**J**) Larval brains were labelled with Fog, Pros and GFP at 24h ALH. White squares, images with higher magnification on the right. White lines, brain lobe outlines. Yellow lines, neuropil outlines. Asterisks, central brain regions. Dash outlines, astrocytes. (**K**) Larval brains were labelled with Fog, Pros at 24h ALH. (**L**), Quantification graph of Fog levels in astrocytes in *control*: 0.64±0.20, n=43; *alrm>Shi^K44A^*: 0.48±0.16, n=22. (**M**)) Larval NSCs (EdU and Dpn) at 24h ALH. (**N**) Quantification graph of EdU-negative NSCs in (M). *alrm*>*control*: 8.3±4, n=12; *alrm*>*Shi^K44A^*: 22.7±5.5, n=10. Yellow arrowheads and dash circles, EdU-negative NSCs. One-Way ANOVA (F) and two-tailed unpaired Student’s t-test (G, H, I, L, N) are used for statistics. ****P<0.0001; **P<0.01; ns, no significance. The means of analyzed phenotypes were shown above each column. Scale bar, 10μm in D, E, J, K and M; 5μm for images with high magnification in J.

We next examined whether Fog plays a role in qNSC reactivation. We knocked down *fog* in glial cells using a pan-glia driver (*repo-GAL4*) and found that the downregulation of Fog in glia caused delayed reactivation of qNSCs (Fig. 7, D and F). By contrast, *fog* knockdown in the NSCs (by *grh*-GAL4 driver) did not affect the reactivation (Fig. S13, A and B). To pinpoint the subtype of glial cells in which Fog mediates qNSC reactivation, various sub-type glial cell drivers were used to specifically knock down *fog* and express mCD8-GFP to label these glial cells: *NP6293-GAL4* and *NP2276-GAL4* drivers for perineurial glia (PG) and subperineurial glia (SPG), respectively; *nrv2-GAL4* driver for cortex glia and ensheathing glia (Fig. S13C), while *alrm-GAL4* with mCD8-GFP specifically decorated astrocytes (Fig. 7C). Remarkably, *fog* knockdown in astrocytes (Fig. 7, E and F; *alrm-GAL4* driver), but not in other glial types (Fig. S13, B and D to F), resulted in defective NSC reactivation. We further marked astrocytes by *alrm>GFP* and nuclear Prospero/Pros. As expected, Fog protein levels in the cell body of astrocytes were significantly reduced and increased upon the knockdown and overexpression, respectively, suggesting an efficient knockdown and Fog antibody specificity (Fig. 7J; Fig. S14; Fig. S15). Fog signal intensity was not obviously altered in the neuropil which contains processes of astrocytes (Fig. 7, I and J; Fig. S14). Perhaps Fog is more stable or relatively resistant to RNAi in these processes, or the Fog signal in the neuropil was non-specific.

Recently, Fog protein has been observed in the sensory neurons and BBB glia of embryonic CNS and at 3rd instar larval brain(*62*), which was inconsistent with its expression pattern in early larval brains. Unexpectedly, Fog protein intensity in the entire central brain region including presumptive BBB glia was significantly reduced upon astrocyte-specific Fog knockdown at 24h ALH (Fig. 7, H and J, asterisks; Fig. S14). Therefore, astrocytes appear to be the major source of Fog in the early larval brain, and Fog protein detected in the central brain including BBB glia may be derived from astrocytes.

Fog protein levels in astrocytes remain similar throughout early brain development (Fig. S16). Moreover, Fog protein levels were not affected by nutritional deprivation (Fig. S16). Therefore, the astrocyte niche appears to be unaffected by the nutritional condition, unlike the known BBB glia niche(*5*).

Fog is known to localize to vesicles derived through dynamin-mediated endocytosis prior to its secretion in the *Drosophila* embryos(*63*). To determine whether astrocytes reactivate qNSCs through dynamin-mediated Fog secretion, we blocked dynamin-mediated endocytosis by overexpressing a dominant negative form of *shibire* K44A (*Shi^K44A^*) in astrocytes using the *alrm-GAL4* driver. Shi^K44A^ overexpression in astrocytes resulted in defective NSC reactivation at 24h ALH (Fig. 7, M and N), suggesting that astrocytes regulate NSC reactivation via Dynamin-mediated Fog secretion. Shi^K44A^ overexpression in the astrocytes strikingly reduced Fog levels in the cell body of astrocytes of the larval brain (Fig. 7, K and L), similar to a previous report on a reduction of Fog protein in the *shibire^[ts1]^* embryos at the restrictive temperature(*63*).

Taken together, Fog protein is produced by astrocytes and functions specifically in these cells to control the NSC exit from the quiescent state.

### Fog promotes quiescent NSC reactivation via Gαq-Rho1-Dia signaling

Next, we tested whether Fog is required for Dia localization in the primary protrusion of qNSCs. Indeed, upon Fog knockdown in astrocytes, Dia protein was significantly reduced in the primary protrusion of qNSCs (Fig. 8, A and B). To determine whether Dia is a physiologically-relevant target of Fog-mediated GPCR signaling, we first tested whether Dia overexpression in NSCs could suppress NSC reactivation defects induced by astrocytes-specific *fog* knockdown. Remarkably, Dia overexpression using a combination of *grh*-Gal4 and *alrm*-Gal4 drivers dramatically suppressed the NSC reactivation defects caused by *fog* knockdown (Fig. 8, C and D), while Dia overexpression using *alrm*-Gal4 driver alone did not (Fig. 8, G and H). This observation suggested that Fog secreted from astrocytes promotes NSC reactivation by regulating Dia localization/function. Likewise, NSC reactivation defects induced by *fog* knockdown were suppressed by overexpression of wild type Gαq or Rho1^GFP^ using a combination of both NSC and astrocyte drivers (Fig. 8 E and F), but not by *alrm -GAL4* driver alone (Fig. 8, G and H). These data indicate that Fog secreted from astrocytes promotes NSC reactivation via Gαq-Rho1-Dia signaling.

**Figure 8.**
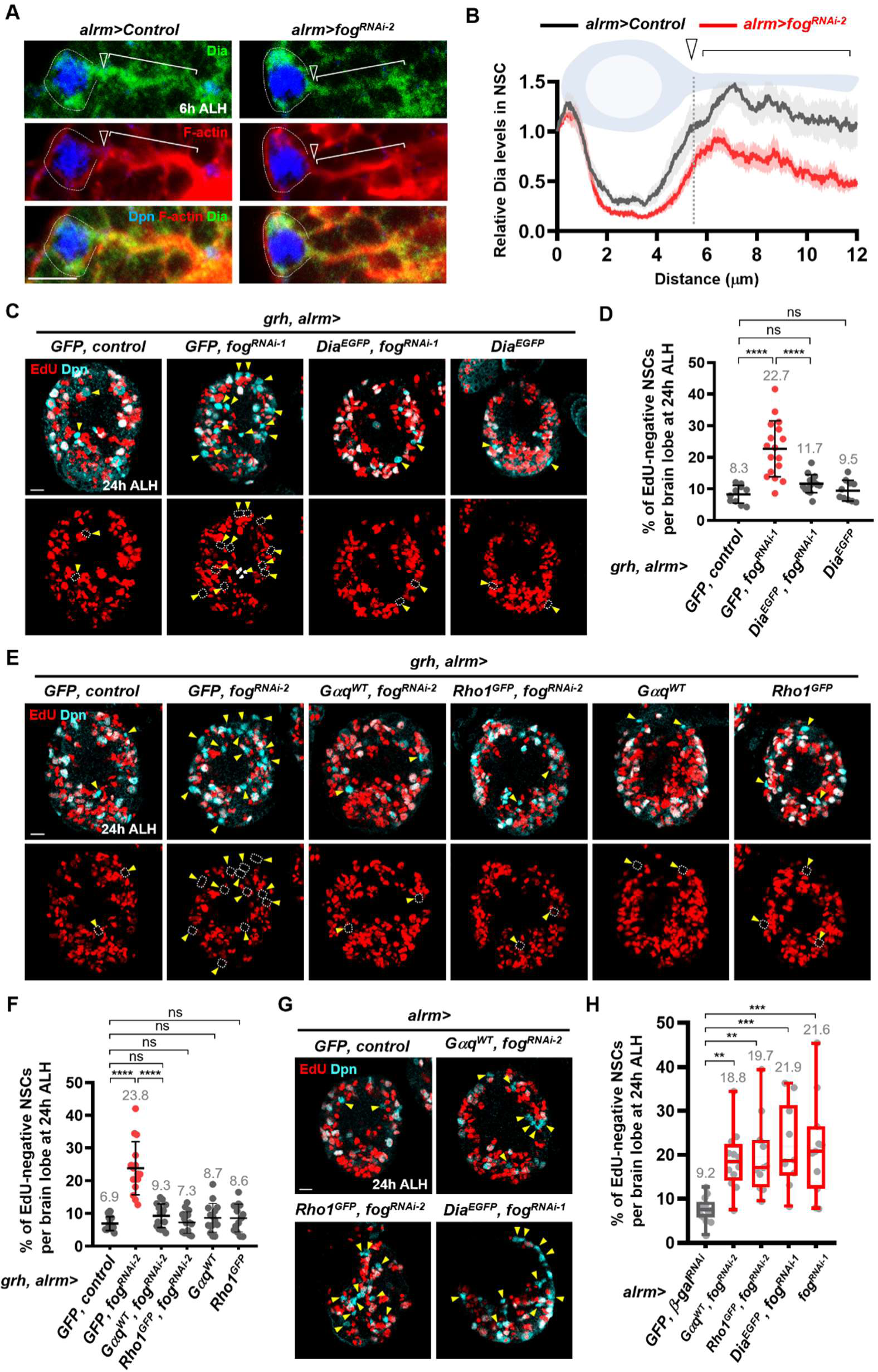
Niche Fog promotes quiescent NSC reactivation via GPCR Smog-Gαq-Dia pathway in NSCs. (**A**) Quiescent NSCs at 6h ALH under the control of *alrm-GAL4* driver were stained for Dia, Dpn and F-actin. White arrowheads, neck region of quiescent NSC. Brackets, primary protrusion marked by F-actin. (**B**) Quantification graph of Dia levels along the soma and protrusion in quiescent NSCs at 6h ALH in control (*β-gal^RNAi^*) and *fog*-KD in astrocyte-like glia. (**C**) Proliferating NSCs (EdU, red; Dpn, cyan) in various genotypes at 24h ALH. (**D**) Quantification graph of EdU-negative NSCs under the control of *grh-, alrm-GAL4* in various genotypes in C. *Control*: 8.3±2.8, n=10; *GFP, fog^RNAi-1^*: 22.7±8.9, n=17; Dia^EGFP^*, fog^RNAi-1^*: 11.7±2.8, n=15; Dia^EGFP^: 9.5±3.3, n=10. (**E**) Proliferating NSCs (EdU, red; Dpn, cyan) at 24h ALH. (**F**) Quantification graph of EdU-negative NSCs under the control of *grh-, alrm-GAL4* in various genotypes in E. *Control*: 6.9±2.1, n=18; *GFP, fog^RNAi-2^*: 23.8±8.1, n=15; *Gαq^WT^, fog^RNAi-2^*: 9.3±3.6, n=16; *Rho1^GFP^, fog^RNAi-2^*: 7.3±3.3, n=15. *Gαq^WT^*: 8.7±4.4, n=15; *Rho1^GFP^*: 8.6±4.3, n=13. (**G**) Proliferating NSCs (EdU, red; Dpn, cyan) in various genotypes at 24h ALH. (**H**) Quantification graph of EdU-negative NSCs under the control of *alrm-GAL4* driver in various genotypes in G. *Control*: 9.2±2.2, n = 8; *Gαq^WT^, fog^RNAi-2^*: 18.8±6.8, n=12; *Rho1^GFP^, fog^RNAi-2^*: 19.7±8.7, n=11; Dia^EGFP^*, fog^RNAi-1^*: 21.9±9.3, n=10; *fog^RNAi-1^*: 21.6±11.2, n=11. One-Way ANOVA was used for statistics. ****P<0.0001; ***P<0.001; **P<0.01; ns, no significance. The means of analyzed phenotypes were showed above each column. Scale bar, 10 μm in C, E and G; 5 μm in A.

## Discussion

In this study, we have uncovered fine F-actin structures and a previously uncharacterized retrograde flow of F-actin in quiescent NSCs by ExM-SIM imaging. Our findings suggest that astrocytes function as a new NSC niche that produce the ligand Fog to activate the GPCR Smog-Gαq-Rho1-Dia signaling; this signaling axis induces F-actin retrograde flow and enhances F-actin polymerization in the soma (Fig. 9, A and B), resulting in the nuclear translocation of Mrtf where it associates with its co-factor SRF/Bs and potentially activates target genes essential for NSC reactivation (Fig. 9C). Mrtf is also required for the expression of the *actin5C* gene (Fig. 9C). Therefore, our work establishes the critical role of the Fog-GPCR Smog-Gαq-Rho1-Diaphanous (Dia)/Formin-MRTF pathway in NSC reactivation.

**Figure 9.**
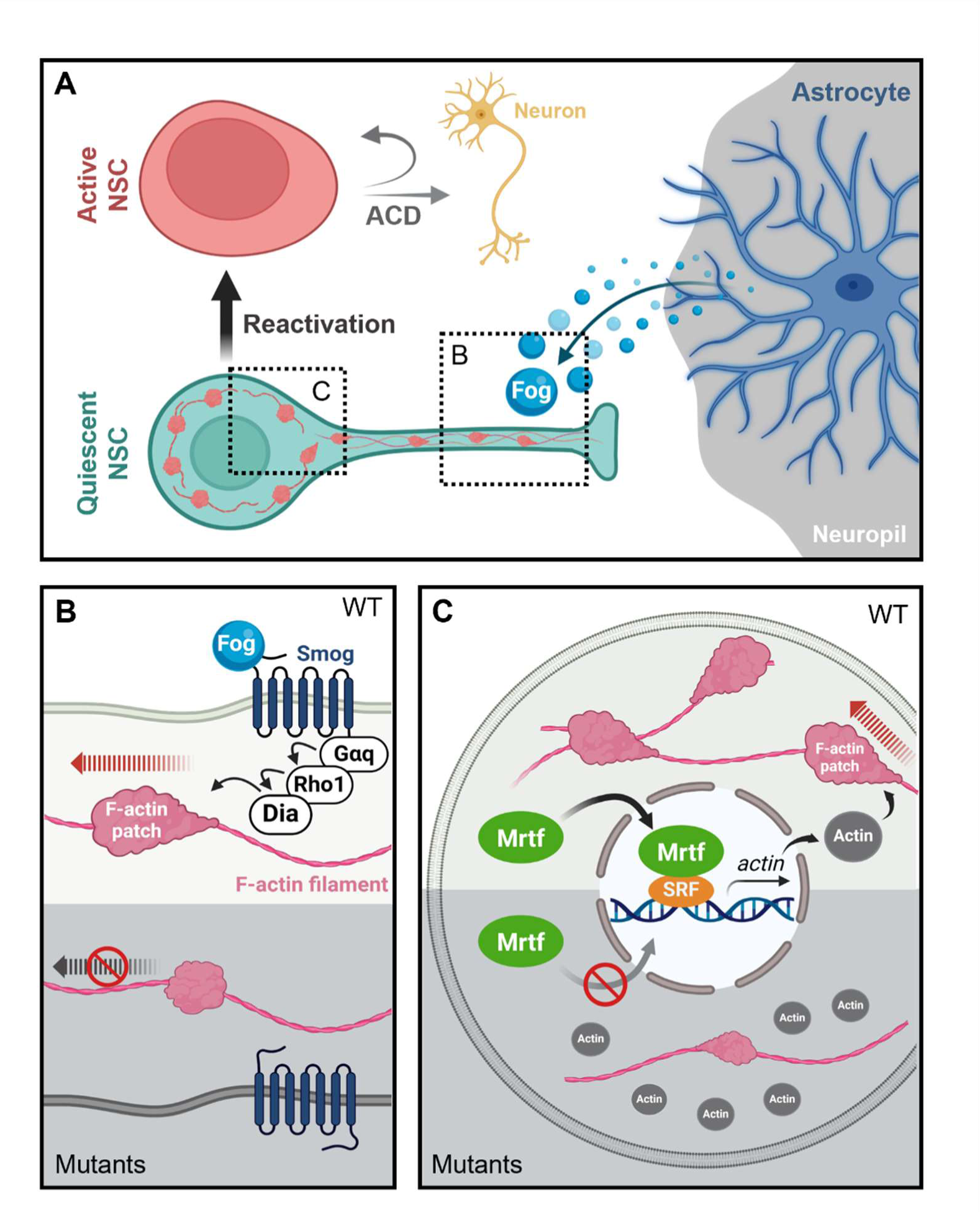
A working model. (**A**) Fog secreted from astrocytes re-activates quiescent NSCs for asymmetric cell division (ACD) of NSC to give rise to new neurons. F-actin forms filaments and patches in quiescent NSCs. (**B**) In the primary protrusion of quiescent NSC, GPCR receptor Smog activated by Fog ligand promotes Gαq-Rho1-Dia signaling in the protrusion, resulting in the retrograde flow of F-actin patches. In the mutants, active Rho1 and Dia cannot transport to the primary protrusion, resulting the reduction of retrograde flow of F-actin patches, leading to the defect of F-actin dynamics in the soma (please see **C**). (**C**) In the soma, F-actin patches from primary protrusion promote robust F-actin polymerization and dynamics to consume G-actin, the monomer of actin. Mrtf can translocate to nucleus and promotes *actin* transcription to feedback to F-actin dynamics and the other unknown target genes that are required for cell proliferation. In the mutants, F-actin amount is reduced, probably due to the defect of retrograde flow of F-actin patches in the primary protrusion, therefore, more G-actin monomers may bind to Mrtf and inhibit Mrtf from nuclear translocation.

### F-actin dynamics in quiescent NSCs

The primary protrusion of quiescent NSCs is enriched with both microtubules and F-actin filaments(*11, 16, 17*). Here, we unravel the structure of these F-actin filaments and characterize their function during quiescent NSC reactivation. We found that F-actin forms filaments and patches in the protrusion of quiescent NSCs, similar to F-actin structures in the axons of cultured mouse hippocampal neurons(*64, 65*). Unlike F-actin multi-filaments formed in the axon shaft(*64, 65*), the majority of protrusions in quiescent NSCs contain two twisted F-actin filaments on which F-actin patches move along. While periodic F-actin rings have been found in the axons and dendrites of neurons using STORM super-resolution microscopy(*64–66*), the protrusion of quiescent NSCs does not seem to have these ring structures. Interestingly, N-methyl-D-aspartate receptor (NMDAR)-mediated Ca^2+^ influx reorganizes F-actin from ring structures to fibers in dendrites(*67*). Quiescent NSC reactivation is synchronized upon nutrient-dependent calcium oscillations of BBB glia in *Drosophila* larval brains(*68*). Since Gαq/PLCβ signaling is known to control calcium dynamics in neural cells(*69*), it would be interesting to understand if calcium signaling regulates F-actin dynamics in quiescent NSC protrusions for NSC reactivation. In quiescent NSCs, the majority of F-actin structures seem to undergo a retrograde flow along F-actin fibers towards the soma. This may re-direct F-actin patches from the protrusion back into the soma for future cell growth and division during reactivation. The retrograde F-actin flow during axonal elongation and guidance is known to facilitate neuronal growth cone motility(*70*). The force of F-actin movement is transmitted to extracellular substrates via cell adhesion molecules on the growth cone(*71*). One such cell adhesion molecule, E-cad, which is required for NSC reactivation was shown to be enriched at the quiescent NSC-neuropil contact sites(*16*). It would be interesting to learn if E-cad at these sites mediates mechanical tension between quiescent NSCs and the neuropil during reactivation.

### Smog/GPR1588-Gαq-Rho1 signaling in brain development and diseases

GPCR signaling regulates a variety of cellular behaviors, including stem cell proliferation and differentiation(*72, 73*). However, the role of GPCR signaling in NSC’s exit from quiescence for reactivation is unknown. We show that the GPCR Smog promotes NSC reactivation via the Gαq-Rho1-Dia pathway. Besides Smog, Mist is another known GPCR for the Fog ligand, and Fog-Mist signaling controls Concertina (Cta, the ortholog of human G Protein Subunit Alpha 13 in *Drosophila*)-Myosin pathway to remodel actin cytoskeleton for apical constriction during embryogenesis(*74–76*). However, knocking down *mist* or *cta* did not cause any NSC reactivation defect (Fig. S17). Furthermore, knocking down *Drosophila* GABA-B-R1 (another GPCR that is highly expressed in NSCs, Fig. S12, A to C) did not affect NSC reactivation (Fig. S17), although mouse GABA-B-R1 is known to couple with Gαq protein to promote calcium influx in neurons during brain development(*77*). These findings validate the specificity of the Smog-Gαq-Dia pathway in quiescent NSCs.

In mammalian systems, the function of Gpr158 (a homologue of *Drosophila* Smog), GNAQ (Gαq homologue), and Rho1/RhoA in neural stem cell proliferation during brain development is not established. Mouse *Gpr158* has been shown to control neural functions to promote hippocampal-dependent memory, spatial learning, and mood control(*58, 78, 79*). Interestingly, Gpr158 interacts with Gαq in the hippocampus where NSCs reside(*58*). Further study is needed to understand whether a conserved Gpr158-GNAQ axis regulates NSC proliferation in the mammalian brain. In humans, activating mutations of Gαq (encoded by the *GNAQ* gene) causes capillary malformations with hyperpigmentation, Sturge-Weber syndrome with brain defects due to hyperproliferation of endothelial cell, and cancers(*80–82*). In the central nervous system, the GTPase RhoA/Rho1 maintains the integrity of adherens junctions by retaining the organization and number of spinal cord neuroepithelium (neural stem cells)(*83*). Consistent with this, inactivating variants of human RHOA cause a neuroectodermal syndrome with linear hypopigmentation(*84, 85*). These reports suggest that Gαq-RhoA signaling may have a general role in promoting cell proliferation during development.

### Dia-Mrtf signaling regulates NSC proliferation and brain development

Variants of MKL2/MRTF are associated with neurodevelopmental disorders including microcephaly and autism spectrum disorder(*55*). Interestingly, we found that *Drosophila Mrtf^KO^*, a null mutant, exhibited a microcephaly-like phenotype (Fig. S18, A and B). However, the function of mammalian MRTF in NSCs or brain development was unknown. Here, we provide strong evidence that Mrtf is a novel intrinsic factor mediating the exit of *Drosophila* NSCs from quiescence. It will be of great interest to understand whether mammalian Mrtf also regulates neural stem cell proliferation. Consistent with the role of *Drosophila* Dia in NSC reactivation, mammalian Formin 2 activates the Wnt signaling-β-Catenin pathway to regulate neural stem cell proliferation during brain development(*86*). Variants of Formins also have been identified in human microcephaly patients(*23, 24, 38*). MRTF is a downstream effector of Formin/Diaphanous-mediated actin dynamics for regulating the transcription of cell motility- and proliferation-related genes in tumor cells and fibroblasts(*87–89*). We found that microcephaly-like phenotype in the larva carrying loss-of-function allele of *dia* can be suppressed by Mrtf overexpression in NSCs, suggesting that Dia-Mrtf signaling plays an important role during brain development.

F-actin dynamics controls cell proliferation via SRF-MRTF signaling in vascular smooth muscle (VSM) cells, where SRF-MRTF signaling activates the transcription of cell proliferation regulators such as Connective tissue growth factor (CTGF) (*90*). We propose that in quiescent NSCs, F-actin dynamics controls NSC reactivation via SRF-MRTF signaling. We speculate that Mrtf might directly target genes that are required for NSC reactivation. In addition, F-actin dynamics might be important for cell growth of quiescent NSCs during their reactivation. *akt*, a component of dInR-PI3K-Akt pathway that controls quiescent NSC reactivation in larval brain of *Drosophila*(*5, 6*), was among the known target genes of Mrtf in *Drosophila* egg chambers(*54*). However, Akt protein levels remained unaltered in *Mrtf*^△^*^7^* NSCs (Fig. S18, C and D). Further studies are required to better understand how F-actin dynamic-mediated Mrtf signaling regulates NSC reactivation during brain development, including identifying its direct targets besides actin.

### Astrocytes function as a new niche to promote quiescence exit of NSCs

Surface glial cells form the BBB in *Drosophila* larval brains and function as an NSC niche to secrete insulin-like peptides that activate quiescent NSCs(*5*). In the mammalian brain, astrocytes contact and surround the BBB, made of endothelial cells, to regulate its permeability; astrocytes also secrete factors including TGF-β, GDNF, and FGF to regulate NSC quiescence, proliferation and differentiation(*91–96*). In this study, we have established that astrocytes, a subtype of glial cells, function as a new NSC niche that promotes NSC reactivation in *Drosophila* larval brains. This astrocyte niche functions independently of the previously known niche, the BBB glial cells. Interestingly, the BBB (surface glial cells) in *Drosophila* is located at the surface of the larval brain, close to the cell body of quiescent NSCs, while astrocytes are located at the neuropil surface, the inner part of the larval brain that comes in contact with the tip of the primary protrusion of quiescent NSCs. Therefore, *Drosophila* quiescent NSCs appear to be interposed between two niches, the BBB glia and astrocytes, for their reactivation. Further, astrocytes produce the ligand Fog, which activates the GPCR Smog signaling pathway in NSCs for their reactivation. As astrocytes-derived Fog signaling is independent of nutritional conditions, Fog/Smog-mediated control of actin dynamics may prime NSCs for reactivation in response to nutritional cues. This study proposes an avenue to manipulate astrocytes and the GPCR signaling pathway to control NSC behaviour for the potential treatment of neurodevelopmental disorders.

## Materials and methods

### Fly stocks and culture

*Drosophila* stocks were cultured at 22–25 °C on standard medium. *yellow white (yw)* and *UAS-β-gal RNAi* were used as wild-type and UAS controls for most experiments. The following fly strains were used in this study: *grh-GAL4* (a gift from Andrea Brand’s Lab), *repo-GAL4* (Bloomington Drosophila stock center (BDSC) #7415), *dia^1^* (BDSC #11762), *dia^5^* (BDSC #9138), *Df(2L)ED1317* (*dia* deficiency, dia^Df^; BDSC #9175), *Gαq^221C^* (BDSC #30744), *Df(2R)Gαq1.3* (*Gαq* deficiency, *Gαq^Df^*, BDSC #44611), *bs[03267]* (BDSC #83157), *Df(2R)Exel6082* (*bs* deficiency, *bs^Df^*, BDSC #7561), *mrtf[Delta7]* (BDSC #58418), *Df(3L)BSC412* (*mrtf* deficiency, *mrtf^Df^* BDSC #24916), Df(2L)Exel9062 (*smog* deficiency, *smog^Df^*, BDSC #7792), *smog knockout(KO)* (*smog^KO^*, a gift from Thomas Lecuit’s’s Lab), *smog*[CR00977-TG4.0] (*smog-GAL4*, BDSC #83229), *sqhP>Smog::GFP* (a gift from Thomas Lecuit’s Lab), *NP6293-GAL4* (Drosophila Genomics Resource Center (DGRC) #105-188), NP2276-GAL4 (DGRC #112-853), *nrv2*-GAL4 (BDSC #6797), *alrm*-GAL4 (BDSC #67032). Flies carrying UAS-RNAi or UAS-transgenes were cultured at 29 °C or 31 °C to enhance the efficiency of gene inhibition or the expression of genes. UAS-RNAi or UAS-transgene lines used in this study are listed as follows: *UAS-GFP-Moesin* (BDSC #31776), *UAS-GFP-UtABD* (a gift from Thomas Lecuit’s Lab), *UAS-β-Gal RNAi* (BDSC #50680), *UAS-dia RNAi-1* (BDSC #33424), *UAS-dia RNAi-2* (BDSC #35479), *UAS-Gαq RNAi-1* (BDSC #33765), *UAS-Gαq RNAi-2* (Vienna Drosophila Resource Center (VDRC) # 105300), *UAS-Gαq^Q203L^* (BDSC #30743), *UAS-rho1 RNAi-1* (BDSC #9909), *UAS-rho1 RNAi-2* (BDSC #9910), *UAS-GFP::Rho1* (BDSC #9393), *UAS-Rho1^N19^* (BDSC #58818), *UAS-DiaRBD-GFP* (BDSC #52291), *UAS-Dia::EGFP* (BDSC #56751), *UAS-bs RNAi-1* (VDRC #330226), *UAS-bs RNAi-2* (BDSC #26755), *UAS-mrtf RNAi-1* (BDSC #31930), *UAS-mrtf RNAi-2* (BDSC #42537), *UAS-mrtf* (BDSC #58421), *UAS-smog RNAi-1* (BDSC #43135), *UAS-smog RNAi-2* (BDSC #51705), *UAS-mist RNAi-1* (BDSC #41930), *UAS-mist RNAi-2* (BDSC #57699), *UAS-cta RNAi-1* (BDSC #41964), *UAS-cta RNAi-2* (BDSC #51849), *UAS-GABA-B-R1 RNAi-1* (BDSC #28353), *UAS-GABA-B-R1 RNAi-2* (BDSC #51817), *UAS-wts RNAi* (BDSC #34064), *UAS-yki-S168A-GFP* (BDSC #28836), *UAS-Gαq^wild type^* (Gαq^WT^, a gift from Wen Hu’s Lab), *UAS-fog RNAi-1* (BDSC #36970), *UAS-fog RNAi-2* (BDSC #61917*), UAS-Shibir^K44A^* (*shi^K44A^*, BDSC #5811), *UAS-spir RNAi-1* (BDSC #61283), *UAS-spir RNAi-2* (BDSC #30516), *UAS-chic RNAi-1* (VDRC #102579), *UAS-chic RNAi-2* (BDSC #34523), *UAS-sn RNAi-1* (BDSC #42615), *UAS-sn RNAi-2* (BDSC #57805), *UAS-Capulet* (BDSC #5943), *UAS-Cpb-mCherry* (BDSC #58727), *UAS-Tsr.N* (BDSC #9234), *UAS-TsrS3A* (BDSC #9236), *UAS-tsr RNAi-1* (VDRC #110599), *UAS-tsr RNAi-2* (BDSC #65055), *UAS-capulet RNAi-1* (BDSC #21995), *UAS-capulet RNAi-2* (BDSC #33010), *UAS-cpb RNAi-1* (BDSC #41952), *UAS-cpb RNAi-2* (BDSC #26298), *UAS-Tsr-S3E* (BDSC #9238), *UAS-Arp2 RNAi-1* (VDRC #29944), *UAS-Arp2 RNAi-2* (VDRC #101999), *UAS-Arp3 RNAi-1* (BDSC #53972), *UAS-Arp3 RNAi-2* (BDSC #32921), *UAS-WASp RNAi-1* (BDSC #25955) and *UAS-WASp RNAi-2* (BDSC #51802).

### Immunohistochemistry

Immunostaining of *Drosophila* larval brain was performed as previously described(*97*). In brief, the brains were dissected in PBS and then fixed in 4% (v/v) formaldehyde /PBS for 22 min at room temperature (RT). Fixed brains were washed with 0.3% PBST (0.3% Triton-X100 in PBS) for three times (10 min each on rotator), blocked with 3% BSA in 0.3% PBST for one hour at RT, and then incubated with primary antibodies in 3% BSA (in 0.3% PBST) overnight at 4 °C. After washing with 0.3% PBST thrice, the brains were then incubated with secondary antibodies in 0.3% PBST for 1.5 hours at RT on a rotator. DNA was labeled with ToPro-3 (1:5000; Invitrogen, Cat #T3605) in 0.3% PBST for 30 min at RT. After washing with 0.3% PBST twice, the brains were mounted in mounting medium. The images were collected on a Zeiss LSM 710 confocal microscope (Axio Observer Z1; ZEISS) using a Plan-Apochromat 40×/1.3 NA oil differential interference contrast objective. They were then processed with Zen software (2010 version), Fuji ImageJ and Adobe Photoshop (V24.1.1) software.

The following primary antibodies were used: guinea pig anti-Dpn (1:1,000), mouse anti-Miranda (Mira, 1:50, F. Matsuzaki), rabbit anti-GFP (1:3,000; F. Yu), mouse anti-GFP (1:5,000; F. Yu), rabbit anti-Dia (1:5,000; S. Wasserman), rabbit anti-RFP (1:2,000), rabbit anti-Yki (1:50), Rabbit anti-Fog antibody (1:200; N.Fuse), rabbit anti-Mrtf antibody (1:200), mouse anti-Repo antibody (1:20; DSHB, Cat #8D12), mouse anti-Prospero antibody (1:10; DSHB, Cat #MR1A), rabbit anti-Akt antibody (1:100, Cell Signaling, Cat #4691S). The secondary antibodies used were conjugated with Alexa Fluor 488, 555 or 647 nms (Jackson laboratory).

Rhodamine-Phalloidin (1:200, Invitrogen, Cat #415) was used to label filamentous actin (F-actin). The larval brains were dissected in PBS and fixed in 4% (v/v) formaldehyde /PBS for 22 min at room temperature (RT), following which the fixed brains were washed with 0.1% PBST (0.1% Triton-X100 in PBS) thrice (10 min each on a rotator). The brains were then blocked with 3% BSA in 0.1% PBST for one hour at RT and incubated with primary antibodies in 3% BSA (in 0.1% PBST) overnight at 4 °C. After washing with 0.1% PBST thrice, the brains were incubated with Rhodamine-Phalloidin and secondary antibodies in 0.1% PBST for 1.5 hours at RT on a rotator. Finally, the brains were mounted in mounting medium. For protein signals measured within the protrusion, the focal plane with the strongest fluorescent signals of the protein of interest was selected for quantification.

### EdU (5-ethynyl-2’-deoxyuridine) incorporation assay

*Drosophila* Larvae were fed with food containing 0.2 mM EdU from Click-iT EdU Imaging Kits (Invitrogen, Cat #C10638) for 4 hours before dissection. The brains were dissected in PBS and fixed with 4% formaldehyde in PBS for 22 min, followed by standard immunohistochemistry. After incubation with secondary antibodies, the brains were washed three times with 0.3% PBST (10 min each), followed by detection of incorporated EdU according to the manufacturers’ protocol.

### Expansion microscopy (EM)-Structure illumination microscopy (SIM)

*Drosophila* larval brains at 6h ALH were dissected in PBS, following which they were immunostained as described above. Poststaining, the brains were incubated in the anchoring solution Acryloyl-X SE (AcX)—0.1 mg/ml AcX in MBS (100 mM MES (2-(N-morpholino) ethanesulfonic acid) and 150 mM NaCl, pH 6.0)—at 4 °C overnight. Next, the samples were incubated in the gelation solution—2 M NaCl, 8.6% sodium acrylate, 2.5% acrylamide, 0.15% bisacrylamide, 0.01% 4-hydroxy-2,2,6,6-tetramenthyl-piperidin-1-oxyl (4-hydroxy-TEMPO), 0.2% tetramethylethylenediamine (TEMED), and 0.2% ammonium persulfate (APS) in PBS at 4°C for three hours, followed by incubation at 37°C for one hour for complete gel polymerization. After gelation, polymerized gels were digested with digestion buffer (8 units/ml proteinase K, 50 mM Tris, pH 8, 1 mM EDTA, 0.5% Triton X-100 in water) on a shaker at room temperature for two hours. Gels were trimmed to small pieces. Small gel pieces were expanded by incubating them in MilliQ water for six hours at room temperature, which increased gel size to about four times their original size. The expanded gels were examined using super-resolution structured illumination microscopy.

The super-resolution spinning disk confocal-structured illumination microscope (SDC-SIM) consisted of a spinning disk platform (Gataca Systems) coupled with an inverted microscope (Nikon Ti2-E; Nikon), a confocal spinning head (CSU-W; Yokogawa), a Plan-Apo objective (100× 1.45-NA), a back-illuminated sCMOS camera (Prime95B; Teledyne Photometrics) and a super-resolution module (Live-SR; GATACA Systems). The system employed a laser combiner (iLAS system; GATACA Systems) that provided excitation light at 488-nm/150 mW (Vortran; for GFP), 561-nm/100 mW (Coherent; for mCherry/mRFP/tagRFP), and 639-nm/150mW (Vortran; for iRFP) wavelengths. All images were acquired and processed using the MetaMorph (Molecular Device) software, followed by further processing with ImageJ and Adobe Photoshop softwares (V24.1.1).

### Live-cell imaging

Larval brains expressing UAS-GFP-Moe or UAS-GFP-utABD under *grh-GAL4* at 6h ALH were dissected in a mixture of Shield and Sang M3 insect medium (Sigma-Aldrich, Cat. S8398) supplemented with 10% FBS. Following dissection, six to eight larval brains were placed in a single well of an 8-chamber borosilicate cover glass containing stabilization medium (0.3% methylcellulose, 10% FBS, 0.05 mg/ml glutathione, and 320 μg/ml insulin in M3 medium). To capture time-lapse images of neural stem cells (NSCs), a super-resolution spinning disk confocal-structured illumination microscopy equipped with a Plan-Apo objective (100× 1.45-NA) was used. The imaging was conducted in a chamber at a temperature of 29°C. Quiescent NSCs with protrusion attached to the neuropil were imaged for 16 hours (5-6 min each time interval). The protrusions of quiescent NSCs were captured with multiple Z-planes to cover the entire thickness of protrusion, thereby preventing any loss of signals resulting from movement out of the focal plane. The videos were processed using Abode Photoshop (V24.1.1) and ImageJ software.

### Fluorescence recovery after photobleaching (FRAP)

Larval brains expressing UAS-GFP::utABD under *grh-GAL4* at 6h ALH were dissected in Shield and Sang M3 insect medium as described in live-cell imaging protocol. FRAP measurements were performed using a laser scanning confocal microscope (40X objective lens and Zoom factor 5) on a Nikon A1R MP. Photo-bleaching was achieved by focusing 25% 488nm laser for 8 secs on the selected region of interest (ROI) in the middle of the protrusion. Fluorescent images of the cells were acquired before and after photobleaching by time-lapse imaging of quiescent NSCs every 1s for 5 mins. The recovery time of fluorescent intensity in ROI of the cell that were photobleached were measured similar to the laser ablation methodology.

### Laser ablation of quiescent NSCs

Larval brains expressing UAS-GFP::utABD under *grh-GAL4* at 6h ALH were dissected in Shield and Sang M3 insect medium (Sigma-Aldrich, Cat. S8398) supplemented with 10% FBS. Dissected brain explants were placed in a well containing M3 medium with 0.3% methylcellulose (see live cell imaging section). Live imaging of larval brains was performed on a Nikon A1R MP laser scanning confocal microscope using 40X objective lens and Zoom factor 5. Quiescent NSCs with protrusion attached to the neuropil were imaged for 15 min (1 min each interval before ablation). The middle region of the NSC protrusions were severed by a pico laser emitting 100-130 nanoWatts (nW) of laser power for 0.5-1 second. After injury, quiescent NSCs were imaged again for 15 min (1 min per interval, 10-15 z-stacks with 0.5-0.8-µm z-intervals). Images were processed and analyzed with ImageJ and Abode Photoshop softwares (V24.1.1).

### Antibody generation

Mrtf antibodies were generated by Abmart (Shanghai, China). The synthetic polypeptides containing the coding sequence of *Drosophila* Mrtf (amino acids 1119–1418) were used as an immunogen to boost rabbits. The antibodies were then subjected to affinity purification to obtain purified polyclonal Mrtf antibodies. Yki antibodies were generated by GeneScript. The synthetic polypeptides containing the coding sequence of *Drosophila* Yki (amino acids 180–418) were used as an immunogen to boost rabbits. The antibodies were then subjected to affinity purification to obtain purified polyclonal Yki antibodies.

### Data analysis of single cell RNA sequencing

Raw data was downloaded from GSE134722 (Brunet *et. al.*, 2019) and processed by Seurat 4.0. The raw data was firstly analyzed according to the methods in Brunet *et. al.* (2019) to separate clusters of neural stem cells, glia and neurons., following which sub-clustering was performed in the clusters of neural stem cells and glial cells. Quiescent and active neural stem cells were annotated by the expression of proliferating markers: *wor*, *CycA*, *CycE*, *PCNA* and etc. Subtypes of glial cells were classified by cell type specific markers: Surface glia: *CG6126* and *Indy*; Cortex/Chiasm glia: *hoe1* and *wrapper*; Astrocyte/Neuropil glia: *wun2*, *Eaat1* and *Gat*. Astrocyte/Neuropil glia were further classified into astrocyte-like glia and ensheathing glia by the astrocyte-like glia specific markers: *e* and *CG31235*.

### Extraction of total messenger RNA (mRNA) and RT-qPCR

Total mRNA was extracted from larval brains of control (yw) and mrtf[Delta7] at 24 h ALH using TRI Reagent (Sigma-Aldrich) according to the manufacturer’s instructions. Reverse transcription was performed with iScript cDNA Synthesis Kit (Bio-RAD) according to the manufacturer’s instructions. RT-qPCR was performed according to the manufacturer’s instructions (SsoFast EvaGreen, Bio-RAD). References genes used as an internal control were as follows: rp49/Rpl32 (Ribosomal protein L32), Sdh (Succinate dehydrogenase), and Tbp1/Rpt5 (Regulatory particle triple-A ATPase 5).

The primers pairs used for RT-qPCR were the following:

rp49 forward: 5’-TGTCCTTCCAGCTTCAAGATGACCATC-3’,
rp49 reverse: 5’-CTTGGGCTTGCGCCATTTGTG-3’;
sdh forward: 5’-GTCTGAAGATGCAGAAGACC-3’,
sdh reverse: 5’-ACAATAGTCATCTGGGCATT-3’.
Tbp-1 forward: 5’-AAGCCCGTGCCCGTATTATG-3’,
Tbp-1 reverse: 5’-AAGTCATCCGTGGATCGGGAC-3’.
mrtf-forward 5’-GAGTCAGCACGTCACTGGAA -3’
mrtf- reverse 5’- ACTCTTTTATGCAGGCGGTG-3’;
actin5C forward 5’- GAGCGCGGTTACTCTTTCAC -3’
actin5C reverse 5’- GCCATCTCCTGCTCAAAGTC -3’.

### Quantification and statistics

*Drosophila* larval brains were placed dorsal side up on microscope slides. Confocal z-stacks were taken from the surface to the deep layers of the larval brains (20–35 z-stacks with 2 or 3 µm intervals per brain lobe). For each genotype, at least 10 brain lobes were collected for z-stack imaging, quantified by ImageJ and plotted in GraphPad Prism 8 software (version 8.3.0). P-values were calculated from two-tailed unpaired Student’s t-test for comparison of two samples. One-way ANOVA, followed by Sidak’s multiple comparisons test was used for comparison of more than two sample groups. All data are shown as the mean ± SD, except for figures 4B, 6H and 8B, which represent mean ± SEM. Statistically-nonsignificant (ns) denotes P > 0.05, * denotes P < 0.05, ** denotes P < 0.01, *** denotes P < 0.001, and **** denotes P < 0.0001. At least two technical replicates were performed for each experiment.

## Acknowledgments

We thank Andrea Brand, Thomas Lecuit, Fumio Matsuzaki, Steven Wasserman, Naoyuki Fuse, Wen Hu, Anuradha Ratnaparkhi, and Fengwei Yu; the Bloomington Drosophila Stock Center, Vienna Drosophila Resource Center, and Kyoto Stock Centre DGGR; the Developmental Studies Hybridoma Bank for fly stocks and antibodies; and Hwei-Jan Hsu for comments on this manuscript. The working model was created with BioRender.

## Funding

This work is supported by the Ministry of Health, Singapore; National Medical Research Council, Singapore MOH-000143 (MOH-OFIRG18may-0004) to H.W.; National Medical Research Council, Singapore Open Fund Young Individual Research Grant MOH-001236 (MOH-OFYIRG22jul-0002) and Khoo Postdoctoral Fellowship award (KPFA/2020/0059) to K.Y.L.; Ministry of Education, Singapore MOE Tier 2 (T2EP30220-0033) to YT; Université de Paris/National University of Singapore grant ANR-18-IDEX-000/2021-03-R/UP-NUS and MBI intramural funding to PK.

## Author contributions

Conceptualization: KYL and HW

Methodology: KYL, MRG, JL, YG, YST, WYD, JH, XT and CSLL

Investigation: KYL, MRG, JL, YG, YST, WYD and JH

Visualization: KYL and HW

Funding acquisition: HW, KYL, PK and YT

Resources: PK and HW

Supervision: HW

Writing—original draft: KYL and HW

## Declaration of interests

The authors declare no competing interests.

## Data and materials availability

All data needed to evaluate the conclusions in the paper are present in the paper and/or the Supplementary Materials.

## Supplementary Materials

### Movie legends

**Movie S1. 3D reconstitution of F-actin structures in qNSCs.** F-actin structures are marked by GFP-Moe, driven by *grh-GAL4*, in qNSCs at 6 h ALH.

**Movie S2. F-actin dynamics in qNSCs.** Top movie, F-actin is marked by GFP-Moe (white), driven by *grh-GAL4*, in qNSCs at 6 h ALH. White arrowheads, retrograde flow of F-actin patches. Scale bar, 5 mm. Bottom movie, thermal theme of F-actin dynamics from top movie.

**Movie S3-1. Injury of protrusion reduces F-actin dynamics in the soma of NSCs.** F-actin is marked by GFP-utABD (black), driven by *grh-GAL4*, in qNSCs at 6 h ALH. White arrowheads, the position of injury caused by laser ablation; black arrows, qNSCs. Scale bar, 5 mm.

**Movie S3-2. Laser ablation in the protrusion of qNSCs.** F-actin is marked by GFP-utABD (black), driven by *grh-GAL4*, in qNSCs at 6 h ALH. Arrows, qNSCs; arrowheads, laser ablation (time 00:06, 00:30 and 00:50). Scale bar, 5 mm. Note that focal planes were changed during the process of laser ablation in the video.

**Movie S4. F-actin dynamics during quiescent NSC reactivation.** F-actin is marked by GFP-utABD (black), driven by *grh-GAL4*, in qNSCs from 6-15 h ALH. Scale bar, 5 mm.

**Movie S5. F-actin retrograde flow is disrupted upon *dia*/*rho1*/*Gαq* depletion.** F-actin is marked by GFP-utABD (white), driven by *grh-GAL4*, in qNSCs of various genotypes at 6 h ALH. White arrows, retrograde flow of F-actin patches. Scale bar, 5 mm.

**Movie S6. Reduced F-actin dynamics in the soma of qNSCs upon *dia*/*rho1*/*Gαq* depletion.** F-actin is marked by GFP-utABD (black), driven by *grh-GAL4*, in qNSCs of various genotypes at 6 h ALH. Scale bar, 5 mm.

**Figure S1.**
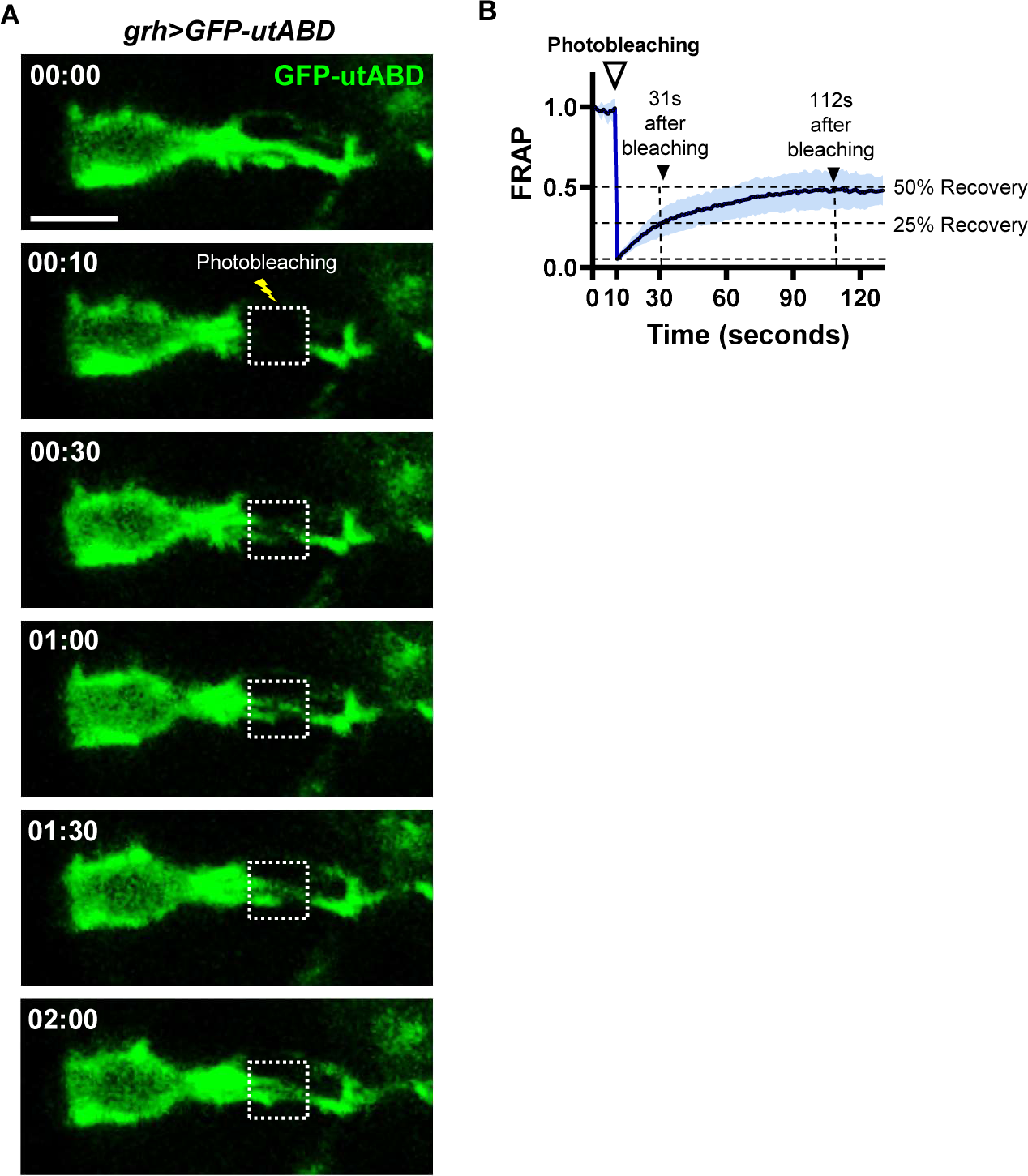
Fluorescence recovery after photobleaching (FRAP) of GFP::utABD in the protrusion of quiescent NSCs at 6h ALH. **A,** Time-lapse images recorded during a fluorescence recovery after photobleaching (FRAP) experiment. Images of GFP-utABD (green) driven by *grh-GAL4* were taken before and after the bleaching. Dash squares indicate bleached area. Time, mm:ss. Scale bar, 5 mm. **B,** Averaged normalized fluorescence recovery of GFP-utABD versus time at bleaching area. White arrowhead indicates the time of photobleaching at 11 seconds. Black arrowheads indicate the time of 25% and 50% recovery at 31 seconds and 112 seconds, respectively. Region of light blue is the standard deviation (S.D.).

**Figure S2.**
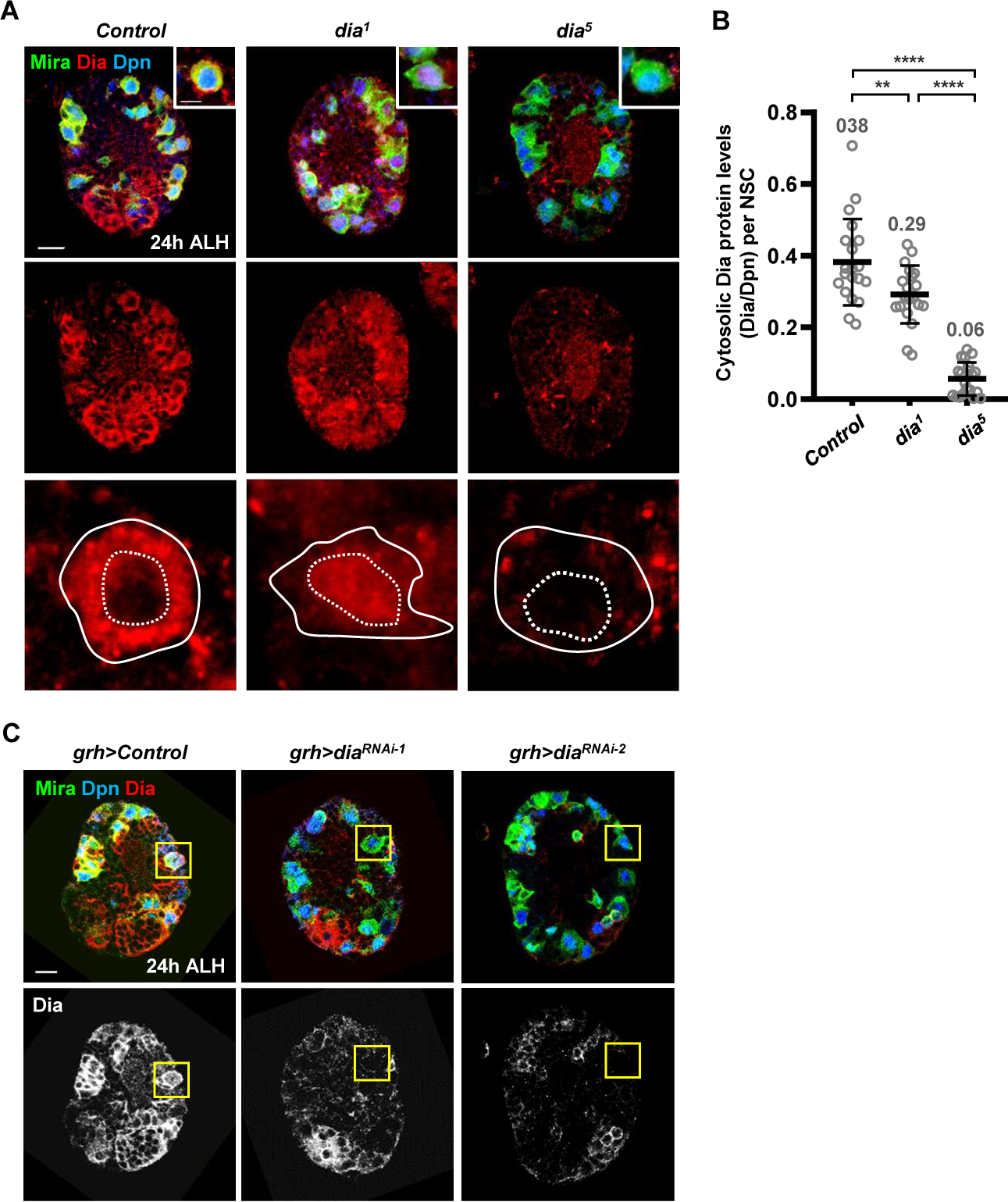
Dia protein is reduced in the *dia*-KD NSCs and NSCs of *dia* mutants. **A,** Dia protein expression (red) in the NSCs (Dpn, blue; Mira, green) in brains of control (*yw*) and *dia* mutants. White squares indicate the region for high magnification. Solid circles, outline of NSC. Dash circles, nucleus. **B,** Quantification of cytoplasmic Dia levels in NSCs of various genotypes in A. **C,** Dia protein expression (red and gray) in the Control and *dia*-KD NSCs (Dpn, blue; Mira, green). Yellow squares indicate NSCs with Dia levels. Scale bar, 10 mm.

**Figure S3.**
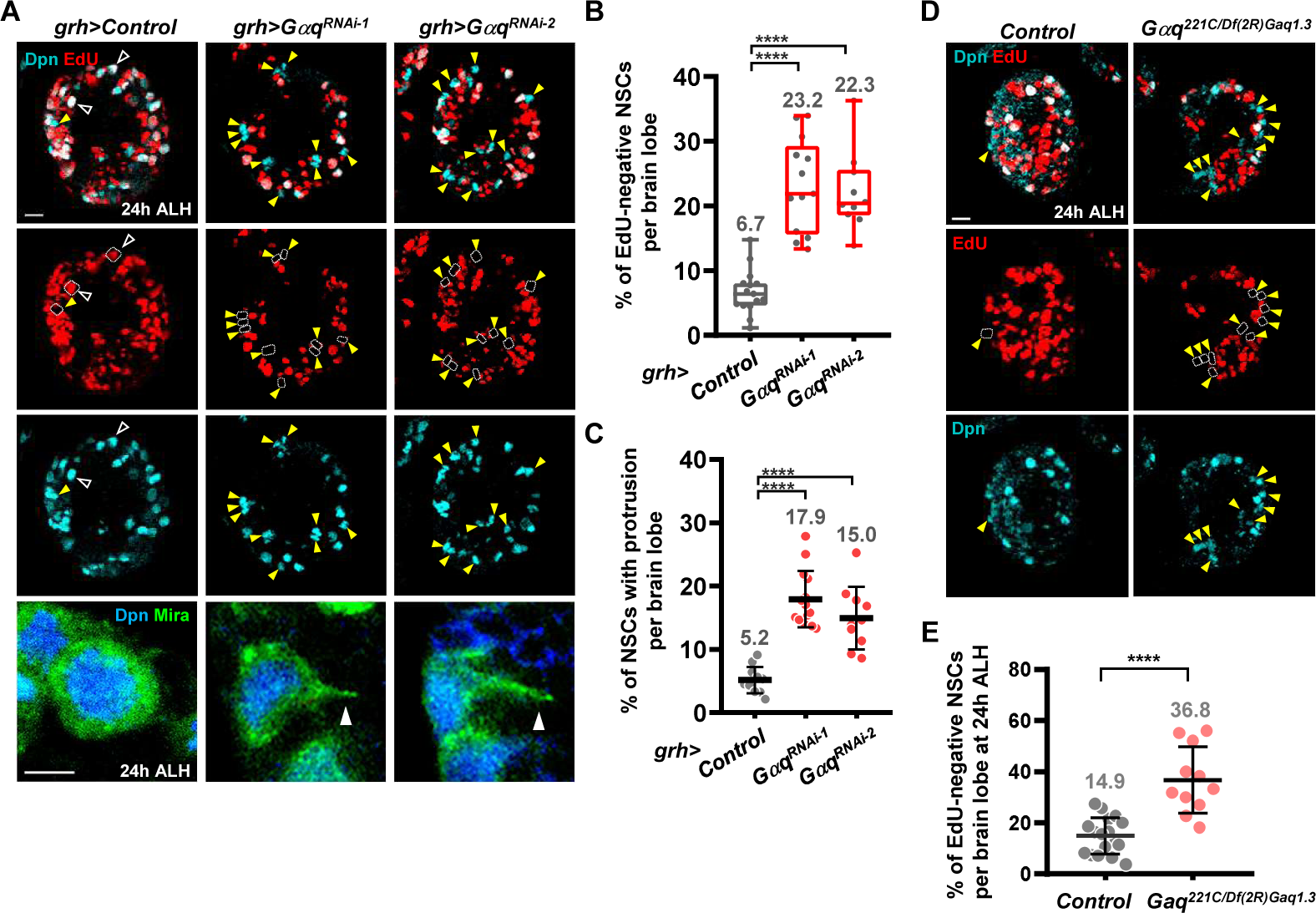
Depletion of Gαq in NSCs causes delayed NSC reactivation. **A,** Proliferating NSCs (EdU, red; Dpn, cyan) in larval brains of control (*β-gal^RNAi^*)- and *Gαq^RNA^*^i^-NSCs at 24h ALH. Yellow arrowheads and dash circles point to EdU-negative quiescent NSCs. White hollow arrowheads point to EdU-positive active NSCs. Scale bar, 10 mm. Bottom row, NSCs (Dpn, blue; Mira, green) in NSCs of control (*β-gal^RNAi^*)- and *Gαq^RNAi^*-NSCs. White hollow arrowheads point to primary protrusion of quiescent NSCs. Scale bar, 5 mm. **B,** Quantification graph of EdU-negative quiescent NSCs in various genotypes in A: Control (*β-gal^RNAi^*): 6.7 ± 3.5, n = 15; *Gαq^RNA^*^i-1^: 23.2 ± 7.2, n = 13; *Gαq^RNA^*^i-2^ : 22.1 ± 6.1, n = 10. **C,** Quantification graph of NSCs retaining primary protrusion in various genotypes in A. Control (*β-gal^RNAi^*): 5.2 ± 2.1, n = 11; *Gαq^RNA^*^i-1^: 17.9 ± 4.5, n = 14; *Gαq^RNA^*^i-2^ : 15.0 ± 5.0, n = 10. **D,** Proliferating NSCs (EdU, red; Dpn, cyan) in larval brains of wild type (*yw*) and *Gαq* mutant larva at 24h ALH. Yellow arrowheads and dash circles point to EdU-negative quiescent NSCs. Scale bar, 10 mm. **E,** The quantification graph of EdU-negative quiescent NSCs in various genotypes. WT: 14.9 ± 7.1, n = 17; *Gαq* mutant: 36.8 ± 13.0, n = 11; One-Way ANOVA (B,C) and *t*-test (E) is used for statistics. ****P < 0.0001. The means of analyzed phenotypes were shown above each column. Box-whiskers plots (B) show the central lines for median values, edges of box represent upper (75th) and lower (25th) quartiles and whiskers show minimum and maximum values.

**Figure S4.**
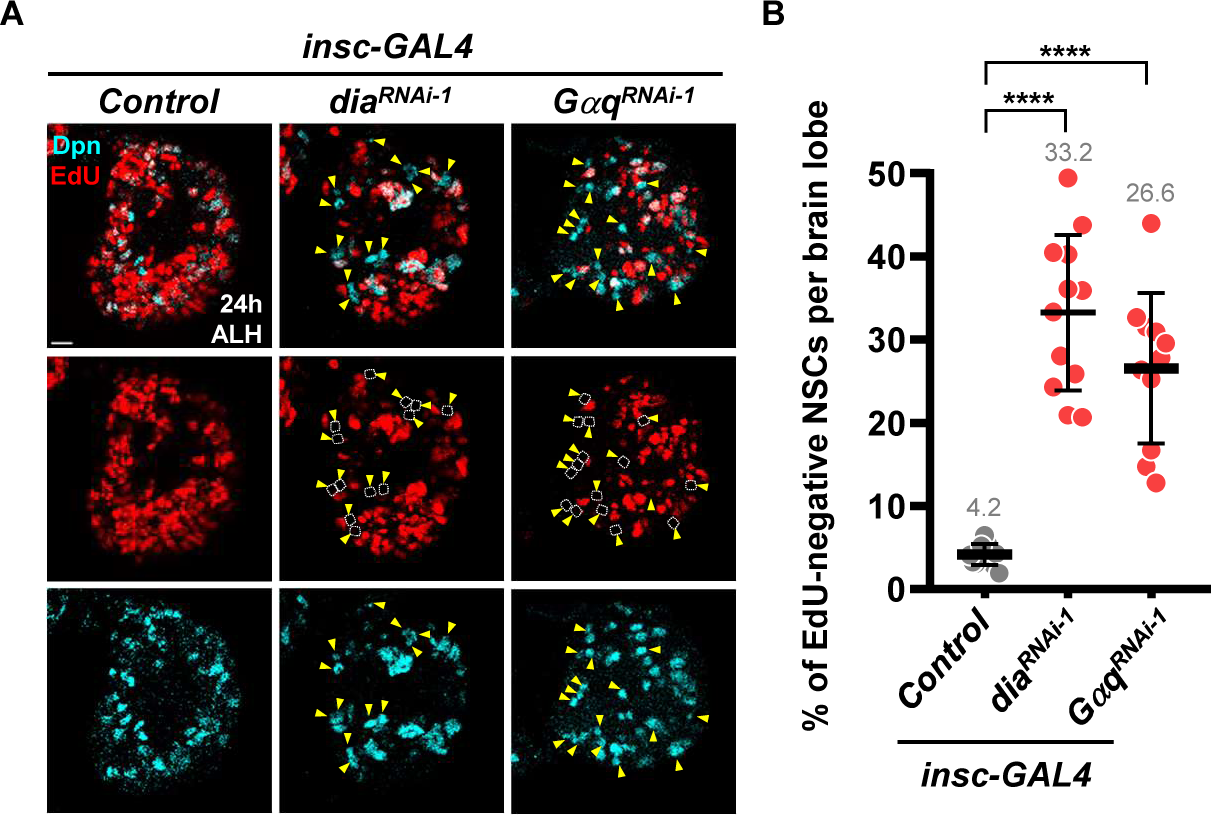
*Gαq* and *dia* knockdown using *insc*-Gal4 results in defects in NSC reactivation. **A,** Proliferating NSCs (EdU, red; Dpn, cyan) in larval brains of various genotypes at 24h ALH. Yellow arrowheads and dash circles point to EdU-negative quiescent NSCs. Scale bar, 10 mm. **B,** Quantification graph of EdU-negative quiescent NSCs in various genotypes in D: Control (*β-gal^RNAi^*): 4.2 ± 1.3, n = 12; *dia^RNAi-1^*: 33.2 ± 9.3, n = 12; *Gαq^RNAi-1^*: 26.6 ± 9.1, n = 11. One-Way ANOVA is used for statistics. ****P < 0.0001. The means of analyzed phenotypes were shown above each column.

**Figure S5.**
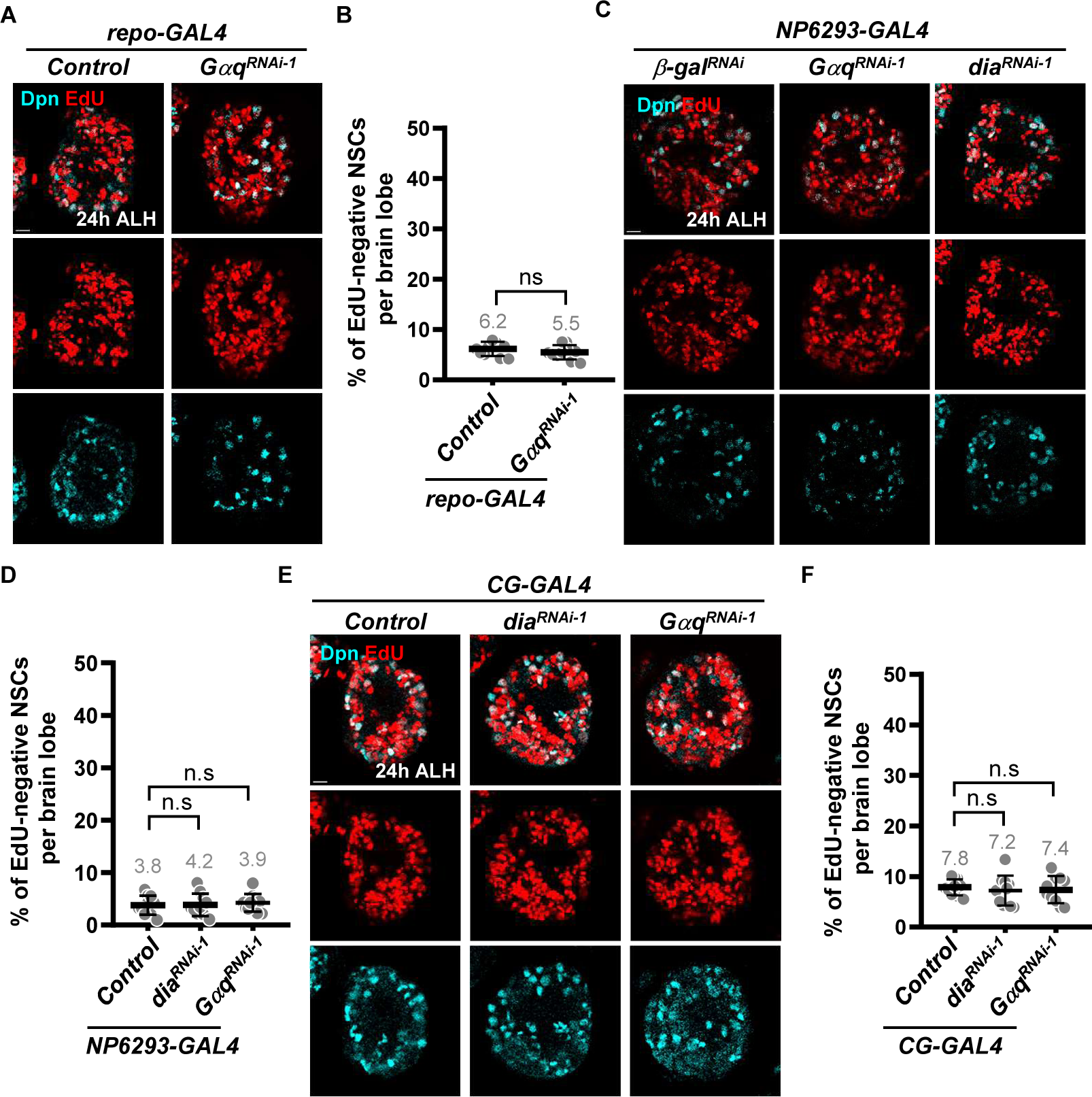
Knockdown *Gαq* or *dia* in glial cells and fat body did not affect NSC reactivation. **A,** Proliferating NSCs (EdU, red; Dpn, cyan) in larval brains carrying pan-glia (*repo-GAL4*) with genes-knockdown at 24h ALH. **B,** Quantification graph of EdU-negative quiescent NSCs in various genotypes in C: Control (*β-gal^RNAi^*): 6.2 ± 1.4, n = 10; *Gαq^RNAi-1^*: 5.5 ± 1.4, n = 10. **C,** Proliferating NSCs (EdU, red; Dpn, cyan) in larval brains carrying BBB glia (*NP6293-GAL4*) with genes-knockdown at 24h ALH. **D,** Quantification graph of EdU-negative quiescent NSCs in various genotypes in C: Control (*β-gal^RNAi^*): 3.8 ± 1.8, n = 10; *dia^RNAi-1^*: 4.2 ± 1.7, n = 10; *Gαq^RNAi-1^*: 3.9 ± 2.1, n = 10. **E,** Proliferating NSCs (EdU, red; Dpn, cyan) of larval brains in larva carrying fat body (*CG-GAL4*) with genes-knockdown at 24h ALH. **F,** Quantification graph of EdU-negative quiescent NSCs in various genotypes in E: Control (*β-gal^RNAi^*): 7.8 ± 1.6, n = 10; *dia^RNAi-1^*: 7.2 ± 2.9, n = 10; *Gαq^RNAi-1^*: 7.4 ± 2.7, n = 10. P-values were calculated from two-tailed unpaired Student’s t-test for comparison of two samples in B and D. One-Way ANOVA is used for statistics in F. ns, no significance. The means of analyzed phenotypes were shown above each column. Scale bar, 10 mm.

**Figure S6.**
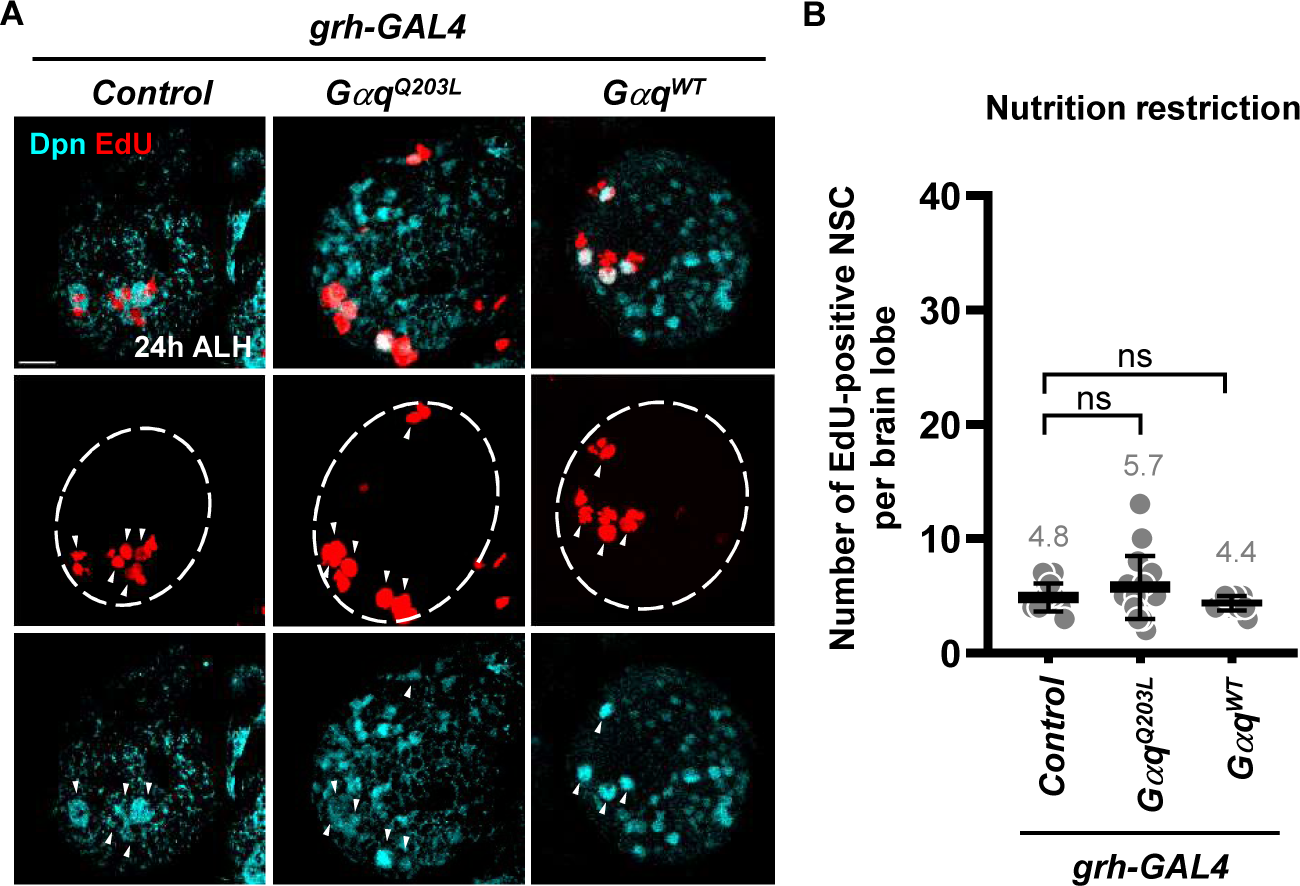
Overexpression of Gαq^Q203L^ and Gαq^WT^ in NSCs did not lead to premature reactivation upon nutrient restriction. **A,** Proliferating NSCs (EdU, red; Dpn, cyan) in larval brains of various genotypes at 24h ALH upon nutrition restriction. **B,** Quantification graph of EdU-positive NSCs in various genotypes in A: Control (*β-gal^RNAi^*): 4.8 ± 1.2, n = 14; *Gαq^Q203L^*: 5.7 ± 2.7, n = 16; *Gαq^WT^*: 4.4 ± 0.6, n = 14. One-Way ANOVA is used for statistics. ns, no significance. The means of analyzed phenotypes were shown above each column. Scale bar, 10 mm.

**Figure S7.**
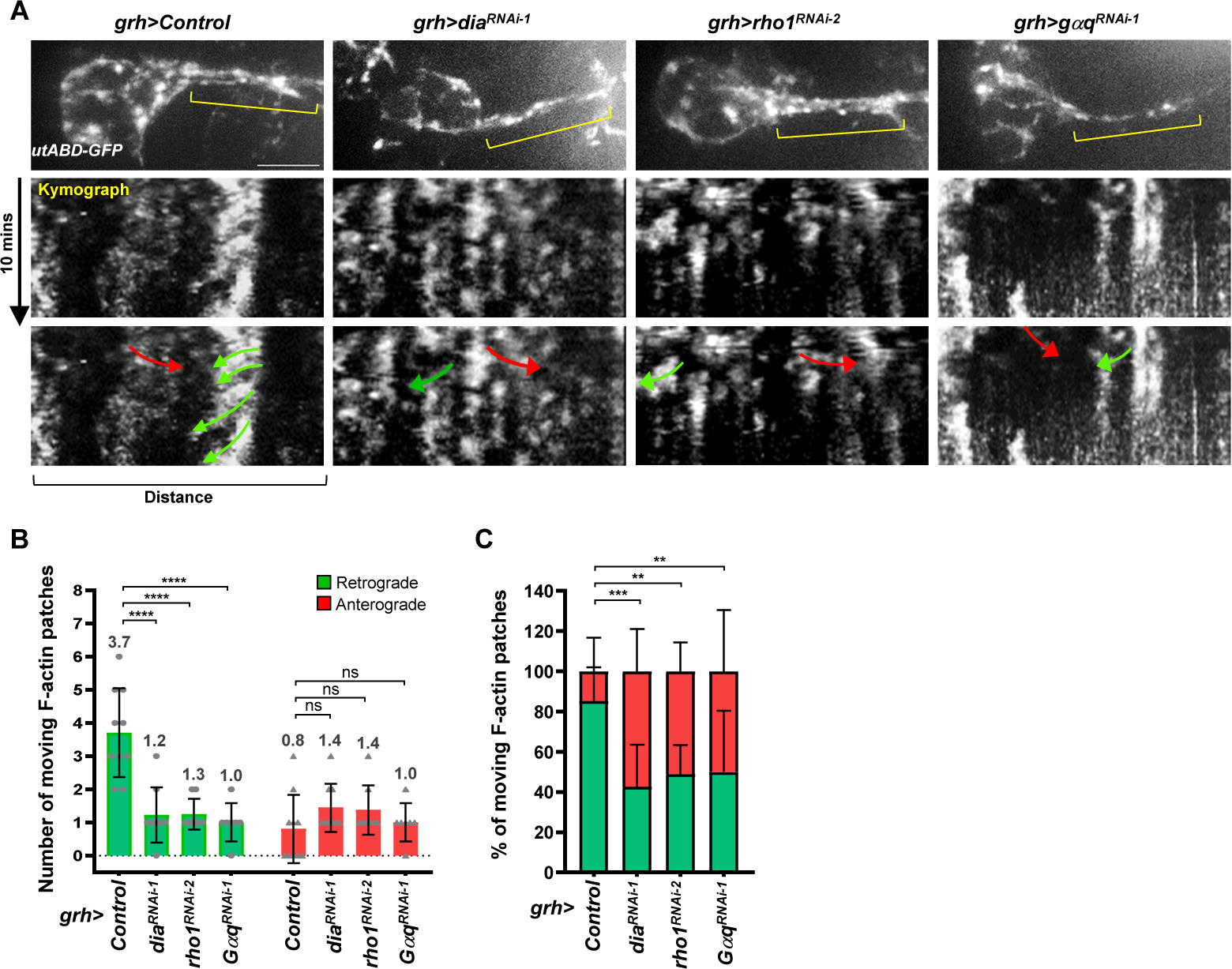
F-actin retrograde flow is impaired in *dia*-, *Gαq*-, *rho1*-KD quiescent NSCs. **A,** Retrograde flow (green arrows) and anterograde flow (red arrows) of F-actin patches in primary protrusions of *dia*-KD, *Gαq*-KD and *rho1*-KD NSCs at 6h ALH. Dash arrows indicate the region of protrusion for kymograph analysis. Total time: 10 mins. Scale bar, 5 mm. **B,** Quantification graph of moving F-actin patches in various genotypes of A. Retrograde: Control (*β-gal^RNAi^*): 3.7 ± 1.3, n = 10; *dia^RNAi-1^*: 1.2 ± 0.8, n = 9; *rho1^RNAi-1^*: 1.2 ± 0.5, n = 8; *Gαq^RNAi^-1*: 1.0 ± 0.6, n = 7. Anterograde: Control (*β-gal^RNAi^*): 0.8 ± 1.0, n = 10; *dia^RNAi-1^*: 1.4 ± 0.7, n = 9; *rho1^RNAi-1^*: 1.4 ± 0.7, n = 8; *Gαq^RNAi-1^*: 1.0 ± 0.6, n = 7. **C,** Quantification graph of percentage of moving F-actin patches in various genetypes. Control (*β-gal^RNAi^*): retrograde 85.2% ± 16.8%, anterograde 14.8% ± 16.8%, n = 10; *dia^RNAi-1^* : retrograde 42.6% ± 21.0%, anterograde 57.4% ± 21.0%, n = 9; ; *Gαq^RNAi-1^*: retrograde 50.0% ± 30.4%, anterograde 50.0% ± 30.4%, n = 7; *rho1^RNAi-1^* : retrograde 48.9% ± 14.3%, anterograde 51.1% ± 14.3%, n = 8. Two-way ANOVA is used for statistics. ****P < 0.0001; ***P < 0.001; **P < 0.01; ns, no significance. The means of analyzed phenotypes were showed above each column.

**Figure S8.**
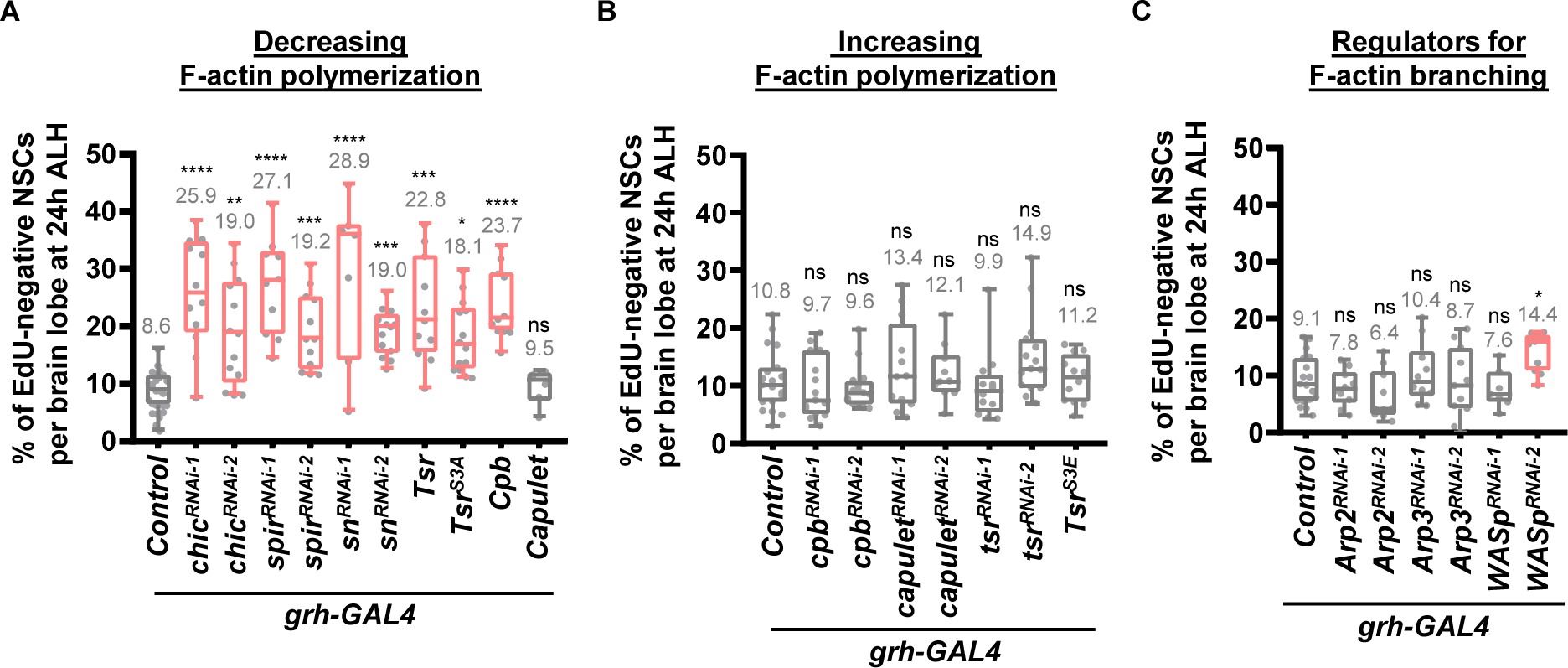
F-actin formation is required for quiescent NSC reactivation. **A,** Quantification graph of EdU-negative quiescent NSCs in various genotypes : Control (*β-gal^RNAi^*): 8.6 ± 3.4, n = 28; *chic^RNAi-1^*: 25.9 ± 9.5, n = 12; *chic^RNAi-2^*: 19.0 ± 9.2, n = 12; *spir^RNAi-1^*: 27.0 ± 8.4, n = 12; *spir^RNAi-2^*: 19.2 ± 6.7, n = 12; *sn^RNAi-1^*: 29.0 ± 14.2, n = 7; sn*^RNA^*^i-2^: 19.0 ± 4.0, n = 13*; Tsr:* 22.8 ± 9.2, n = 11; *Tsr^S3A^*: 18.1 ± 6.2, n = 14; *Cpb:* 23.7 ± 6.1, n = 11; *Capulet*: 9.5 ± 3.0, n = 6. **B,** Quantification graph of EdU-negative quiescent NSCs in various genotypes : Control (*β-gal^RNAi^*): 10.8 ± 5.2, n = 19; *cpb^RNAi-1^*: 9.7 ± 5.4, n = 15; *cpb^RNAi-2^*: 9.6 ± 3.7, n = 15; *Capulet^RNAi-1^*: 13.4 ± 7.8, n = 12; *Capulet^RNAi-1 RNA^*^i-2^: 12.1 ± 5.0, n = 10; *tsr^RNAi-1^*: 9.9 ± 6.1, n = 12; *tsr^RNAi-2^*: 15.0 ± 7.0, n = 15; *Tsr^S3E^*: 11.2 ± 4.2, n = 12. **C,** Quantification graph of EdU-negative quiescent NSCs in various genotypes : Control (*β-gal^RNAi^*): 9.1 ± 4.3, n = 18; Arp2*^RNAi-1^*: 7.8 ± 3.2, n = 10; Arp2*^RNAi-2^*: 6.4 ± 4.3, n = 10; Arp3*^RNAi-1^* : 10.4 ± 5.3, n = 10; Arp3*^RNAi-2^* :8.7 ± 6.2, n = 11; *WASp^RNAi-1^*: 7.6 ± 3.4, n = 7; *WASp^RNAi-2^* : 14.4 ± 3.3, n = 11. One-Way ANOVA is used for statistics. ****P < 0.0001; ***P < 0.001; **P < 0.01. *P < 0.05; ns, no significance. The means of analyzed phenotypes were shown above each column. Box-whiskers plots (A, B and C) show the central lines for median values, edges of box represent upper (75th) and lower (25th) quartiles and whiskers show minimum and maximum values.

**Figure S9.**
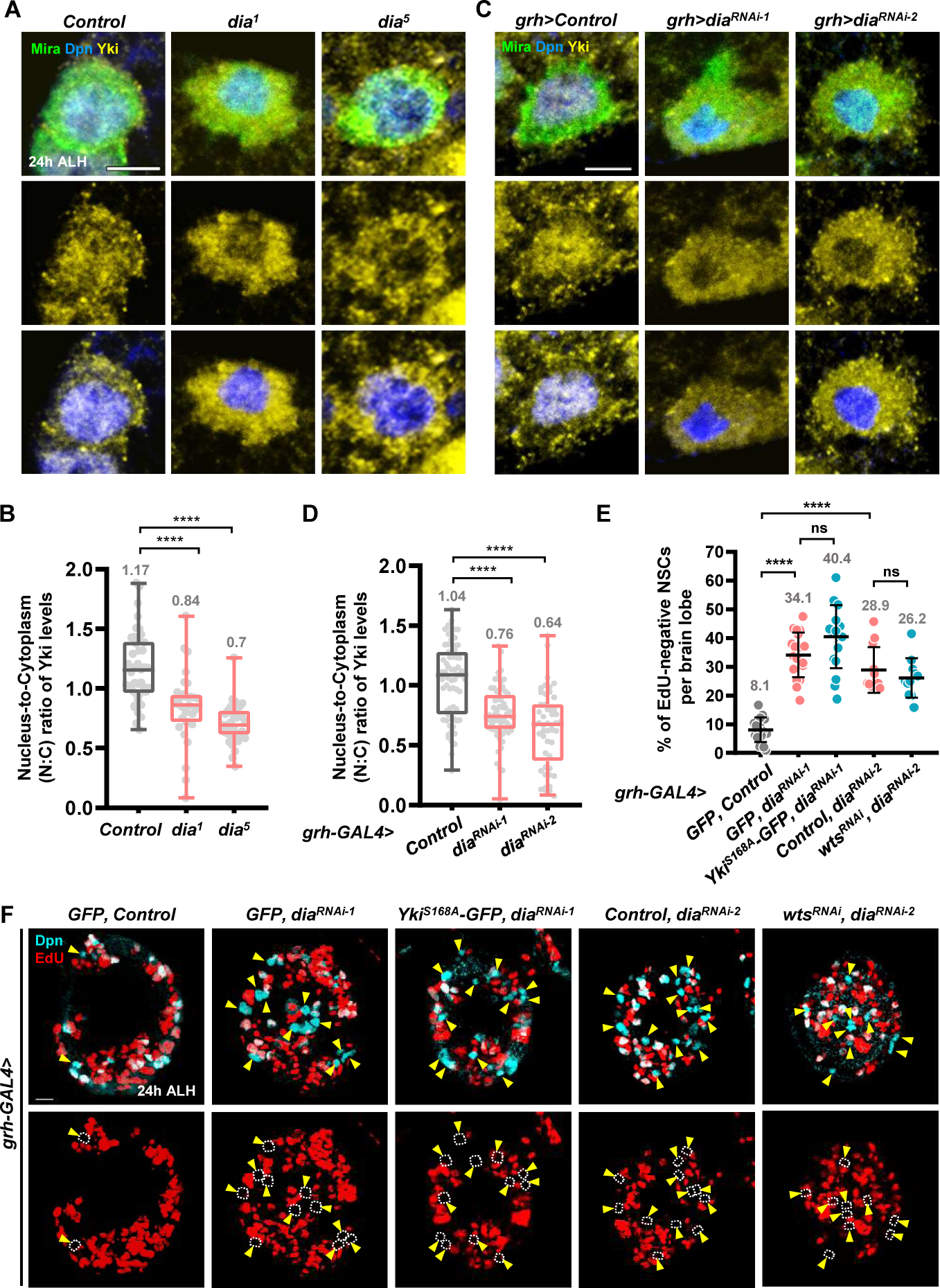
Dia-mediated NSC reactivation is not through Hippo-Wts-Yki pathway. **A,** Yki (yellow) levels in the NSC (Dpn, blue; Mira, green) of control(*yw*) and *dia* mutants larval brain at 24h ALH. **B,** Quantification of nucleus-to-Cytoplasm (N:C) ratio of Yki levels in NSCs of various genotypes of A. **C,** Yki (yellow) levels in the NSC (Dpn, blue; Mira, green) of Control(b-gal^RNAi^) and *dia^RNAi^* lines at 24h ALH. **D,** Quantification of nucleus-to-Cytoplasm (N:C) ratio of Yki levels in NSCs of various genotypes of C. **E,** Quantification graph of EdU-negative quiescent NSCs in various genotypes in A: Control (*β-gal^RNAi^*), GFP: 8.1 ± 4.3, n = 16; *dia^RNAi-1^*, GFP: 34.1 ± 7.8, n = 19; *dia^RNAi-1^, Yki^S168A^-GFP*: 40.4 ± 11.0, n = 18; *dia^RNAi-2^, GFP*: 28.9 ± 7.9, n = 12; *dia^RNAi-2^, wts^RNAi^* : 26.2 ± 7.9, n = 11. **F,** Proliferating NSCs (EdU, red; Dpn, cyan) in larval brains of various genotypes at 24h ALH. Yellow arrowheads and dash circles point to EdU-negative quiescent NSCs. One-Way ANOVA is used for statistics. ****P < 0.0001; ns, no significance. The means of analyzed phenotypes were shown above each column. Scale bar, 5 mm for A and C, 10 mm for F.

**Figure S10.**
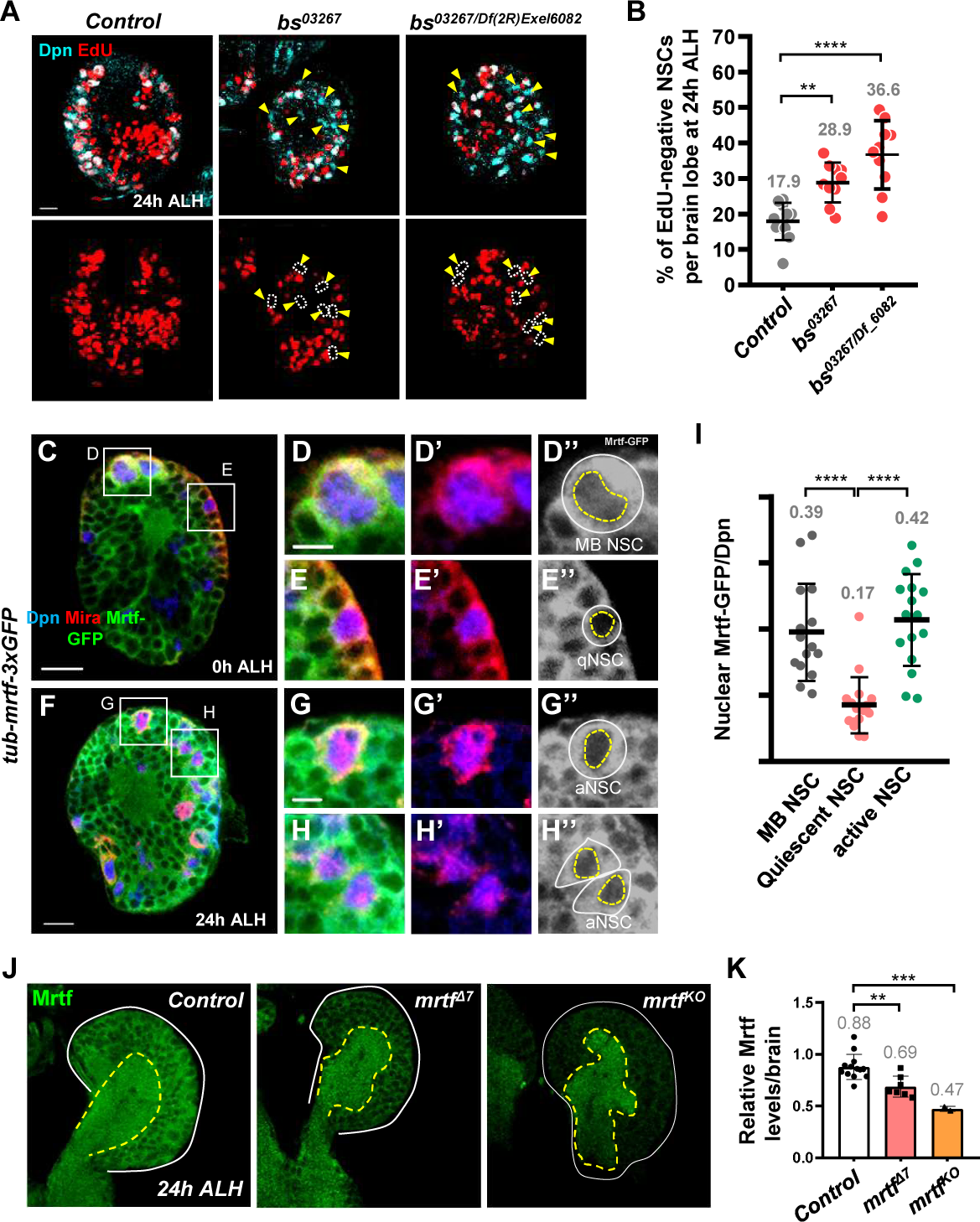
*bs* is required for NSC reactivation and nuclear Mrtf correlates with active status of NSC in larval brains. **A,** Proliferating NSCs (EdU, red; Dpn, cyan) in larval brains of wild type (*yw*) and *bs* mutants at 24h ALH. Yellow arrowheads and dash circles point to EdU-negative quiescent NSCs. Scale bar, 10 mm. **B,** Quantification graph of EdU-negative quiescent NSCs in various genotypes in A: Wild type (*yw*): 17.9 ± 5.2, n = 11; *bs^03267^*: 28.9 ± 5.6, n = 10; *bs^03267/Df(2R)Exel6082^* : 36.6 ± 9.6, n = 10. **C-H,** Mrtf-GFP (green and gray) in NSCs (Dpn, blue; Mira, red). Mushroom body NSCs (MB NSCs) and quiescent NSCs (qNSCs) of brain at 0h ALH and active NSCs (aNSCs) of brain at 24h ALH. White outlines, the edge of cells; yellow dash lines, the nucleus. **I,** Quantification graph of Mrtf-GFP levels among different types of NSCs. **J,** Mrtf levels in various *Mrtf* mutants. White lines, edge of brains. Dash outlines, neuropil region. **K,** Quantification graph of Mrtf levels of brains in various genotypes in E. **L,** Quantification graph of RT-qPCR analysis in 24 h ALH brains from control (*yw*) and *mrtf^Δ7^*. After normalization against *yw* control (with SD):, *mrtf*, 0.52 ±0.08-fold; *actin5C*, 0.55 ±0.03-fold; *Hsp23*, 1.00 ±0.11-fold. One-Way ANOVA is used for statistics. ***P < 0.001; ns, no significance. The means of analyzed phenotypes were shown above each column.

**Figure S11.**
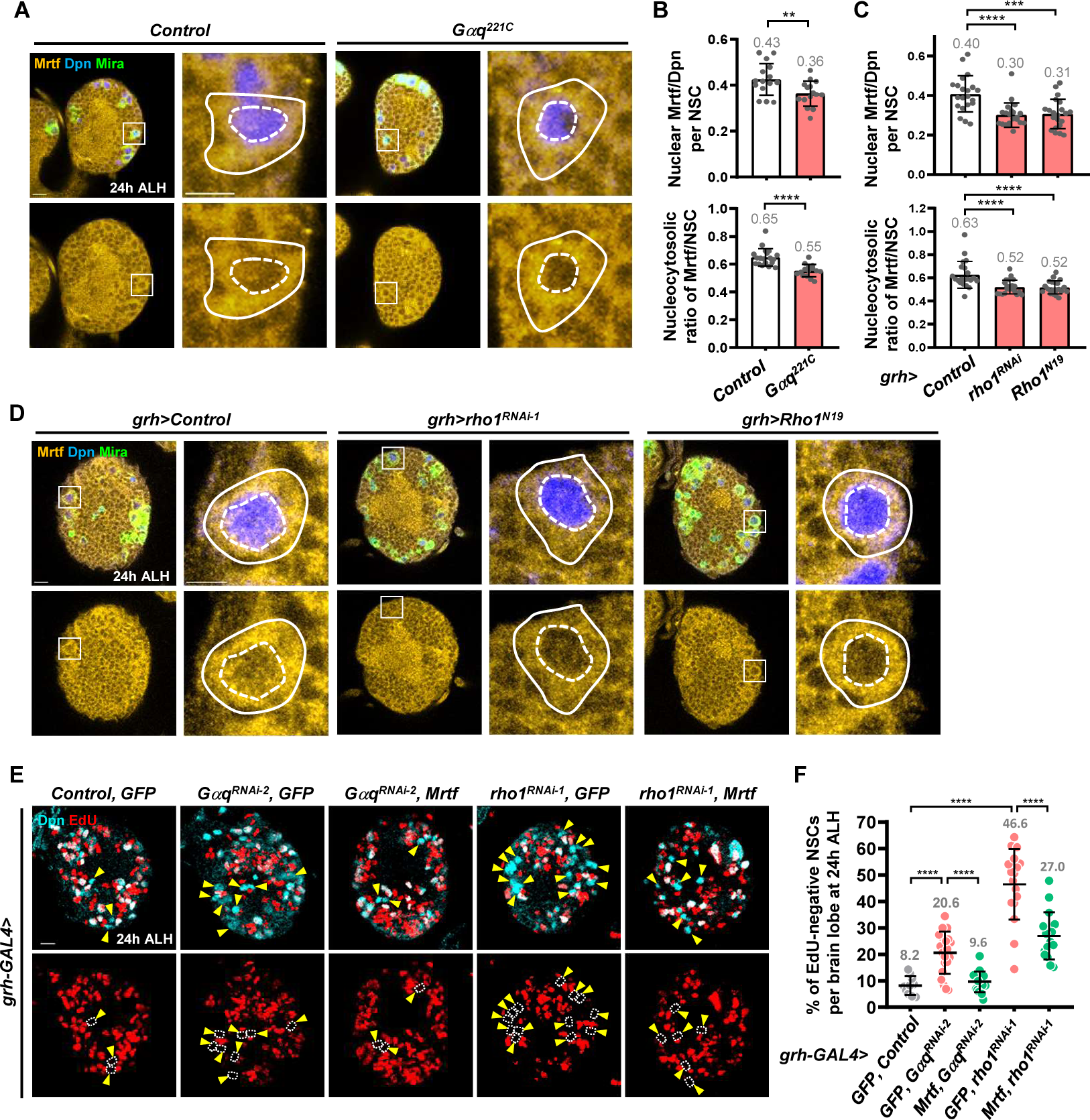
Gαq-Rho1 pathway promotes quiescent NSC reactivation via Mrtf. **A,** Mrtf protein expression (yellow) in the NSCs (Dpn, blue; Mira, green) of different genotypes. White squares, the region for high magnification. Solid outlines, outline of NSC. Dash outlins, nucleus. Scale bar, 10 mm; 5 mm for image with high magnification. **B,** Quantification graph of nuclear Mrtf expression (top) and ratio of nuclear Mrtf to cytoplasmic Mrtf (bottom) in control (*yw*), dia5 and mrtfD7 mutants. Top graph: *control* (*yw*): 0.66 ± 0.2, n = 16; *dia^5^*: 0.44 ± 0.14, n = 15; *mrtf*^D^*^7^*: 0.46 ± 0.14, n = 16. Bottom graph: *Control* (*yw*): 0.71 ± 0.06, n = 16; *dia^5^*: 0.59 ± 0.07, n = 15; *mrtf*^D^*^7^*: 0.56 ± 0.06, n = 19. **C,** Quantification graph of nuclear Mrtf (top) and ratio of nuclear Mrtf to cytoplasmic Mrtf (bottom) in wild Control (*β-gal^RNAi^*),. Top graph: *Control* (*yw*): 0.66 ± 0.2, n = 16; *dia^5^*: 0.44 ± 0.14, n = 15; *mrtf*^D^*^7^*: 0.46 ± 0.14, n = 16. Bottom graph: control (*yw*): 0.71 ± 0.06, n = 16; *dia^5^* : 0.59 ± 0.07, n = 15; *mrtf*^D^*^7^*: 0.56 ± 0.06, n = 19. **D,** Mrtf protein (yellow) in the NSCs (Dpn, blue; Mira, green) of different genotypes. White squares circle the region for high magnification. Solid circles, outline of NSC. Dash circles, nucleus. Scale bar, 10 mm; 5 mm for image with high magnification. **E,** Proliferating NSCs (EdU, red; Dpn, cyan) in larval brains of various genotypes at 24h ALH. Yellow arrowheads and dash circles point to EdU-negative quiescent NSCs. Scale bar, 10 mm. **F,** Quantification graph of EdU-negative quiescent NSCs in various genotypes in A: Control (*β-gal^RNAi^*), GFP: 8.2 ± 3.5, n = 10; *Gαq^RNA^*^i-2^, GFP: 20.6 ± 8.0, n = 21; *Gαq^RNA^*^i-2^, Mrtf: 9.6 ± 3.9, n = 17; *rho1^RNA^*^i-1^, GFP: 46.5 ± 13.3, n = 17; *rho1^RNA^*^i-1^, Mrtf: 27.0 ± 8.9, n = 16. One-Way ANOVA is used for statistics. ****P < 0.0001; ***P < 0.001; **P < 0.01. The means of analyzed phenotypes were shown above each column.

**Figure S12.**
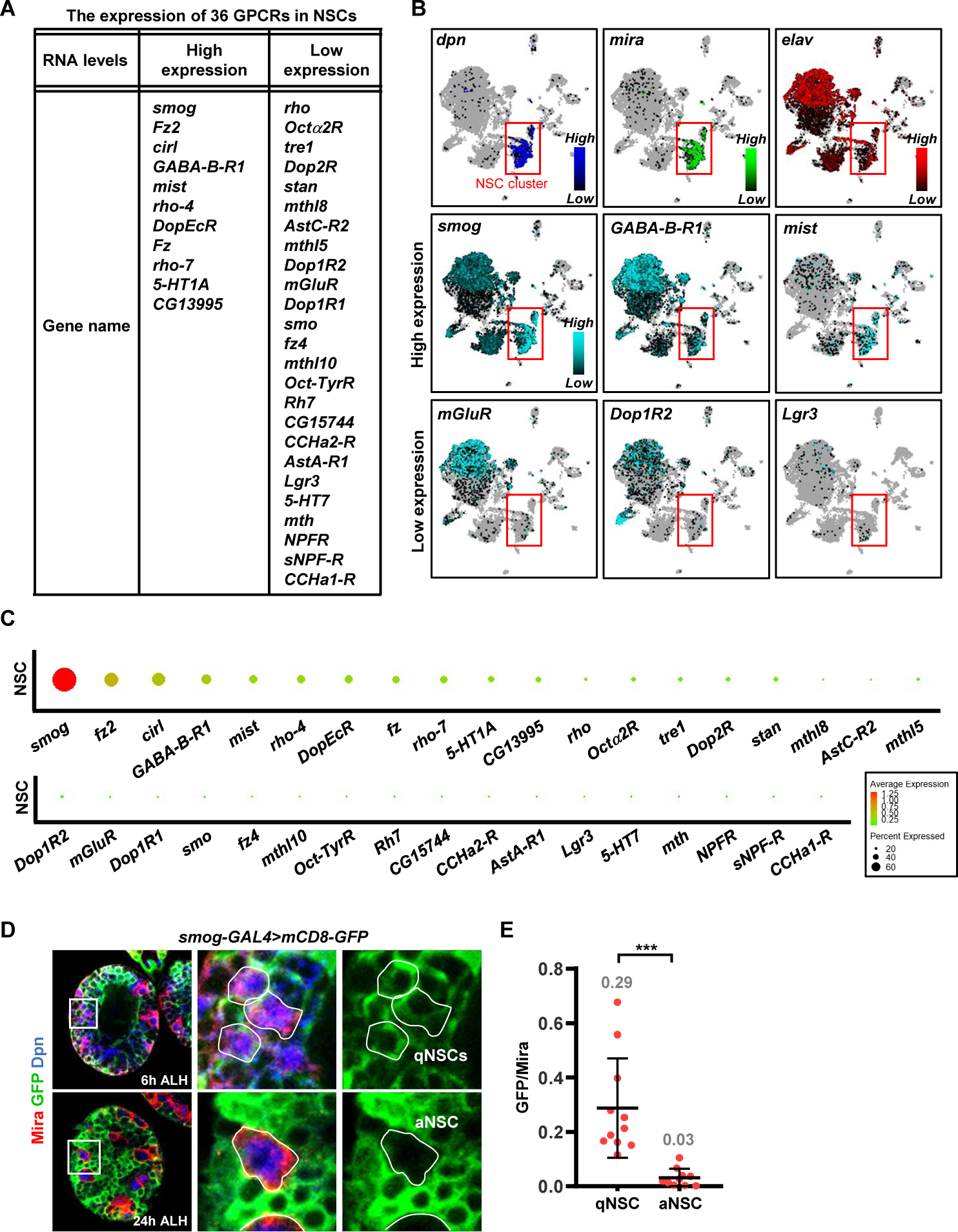
*smog* expression is higher in the quiescent NSCs than active NSCs. **A,** The table of the expression of 36 GPCRs in NSCs. **B,** The t-distributed stochastic neighbor embedding (t-SNE) maps for larval brains at 16h ALH from SCope. NSC markers (*dpn* and *mira*), neural marker (*elav*) and represented high and low GPCRs expression (Cyan). Gradient bars inside the box indicate high to low expression of individual gene in a single cell within a t-SNE map. **C,** Analysis of expression levels of 36 GPCRs in NSCs of larval brain at 16h ALH (dataset of single cell RNA seq from Brunet Avalos, C. et al., 2019). **D,** The transcriptional activity of GPCR *smog* shown by *smog-GAL4>mCD8-GFP* in brains of 6h ALH larva and 24h ALH larva. Quiescent NSCs, qNSCs; active NSCs, aNSCs. White lines mark the edge of indicative cells. **E.** Quantification of GFP levels in quiescent NSCs (qNSCs) and active NCSs (aNSCs). qNSCs: 0.29 ± 1.8, n = 11; aNSCs: 0.03 ± 0.03, n = 10. The means of analyzed phenotypes were shown above each column. P-values were calculated from two-tailed unpaired Student’s t-test for comparison of two samples.

**Figure S13.**
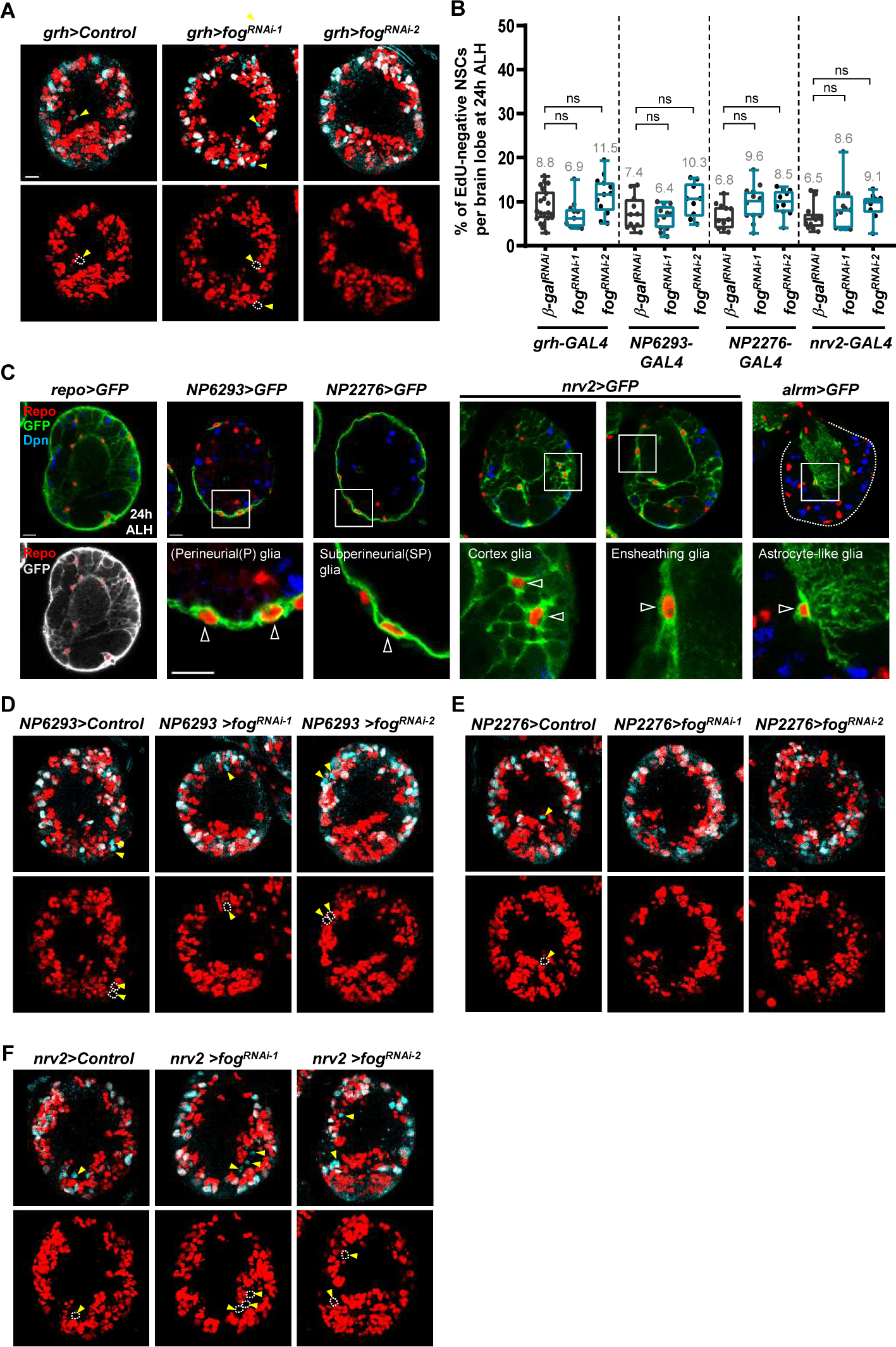
Fog is not required in NSCs, surface glia, ensheathing glia or cortex glia during quiescent NSC reactivation. **A,** Proliferating NSCs (EdU, red; Dpn, cyan) in larval brains carrying *fog*-KD NSCs. Yellow arrowheads and dash circles point to EdU-negative quiescent NSCs. White hollow arrowheads point to EdU-positive active NSCs. **B,** Quantification graph of EdU-negative quiescent NSCs in various genotypes : *grh>Control(β-gal^RNAi^*): 8.8 ± 3.8, n = 21; *grh>fog^RNAi-1^*: 6.9 ± 3.3, n = 11; *grh>fog^RNAi-2^*: 11.5 ± 4.3, n = 12; *NP6293>Control(β-gal^RNAi^*): 7.4 ± 3.8, n = 11; *NP6293>fog^RNAi-1^*: 6.4 ± 2.7, n = 12; *NP6293>fog^RNAi-2^*: 10.3 ± 3.8, n = 9; *NP2276>Control(β-gal^RNAi^*): 6.8 ± 2.8, n = 11; *NP2276>fog^RNAi-1^*: 9.6 ± 4.1, n = 12; *NP2276>fog^RNAi-2^*: 9.8 ± 2.9, n = 10; *nrv2>Control(β-gal^RNAi^*): 6.5 ± 2.9, n = 12; *nrv2>fog^RNAi-1^*: 8.6 ± 4.9, n = 13; *nrv2>fog^RNAi-2^*: 9.1 ± 2.9, n = 9. One-Way ANOVA is used for statistics. ns, no significance. **C,** The patterns of UAS-mCD8-GFP driven by glial type-specific GAL4. Repo (red), pan-glial marker. Dpn (blue), neural stem cell marker. GFP (green and gray), White triangles, glial types under different glial GAL4 lines. **D-F,** Proliferating NSCs (EdU, red; Dpn, cyan) in larval brains carrying *fog*-KD perineurial glial cells (D), *fog*-KD subperineurial glial cells (E) or *fog*-KD cortex/ensheathing glial cells (F) at 24h ALH. Scale bar, 10 mm. The means of analyzed phenotypes were shown above each column. Box-whiskers plots (B) show the central lines for median values, edges of box represent upper (75th) and lower (25th) quartiles and whiskers show minimum and maximum values.

**Figure S14.**
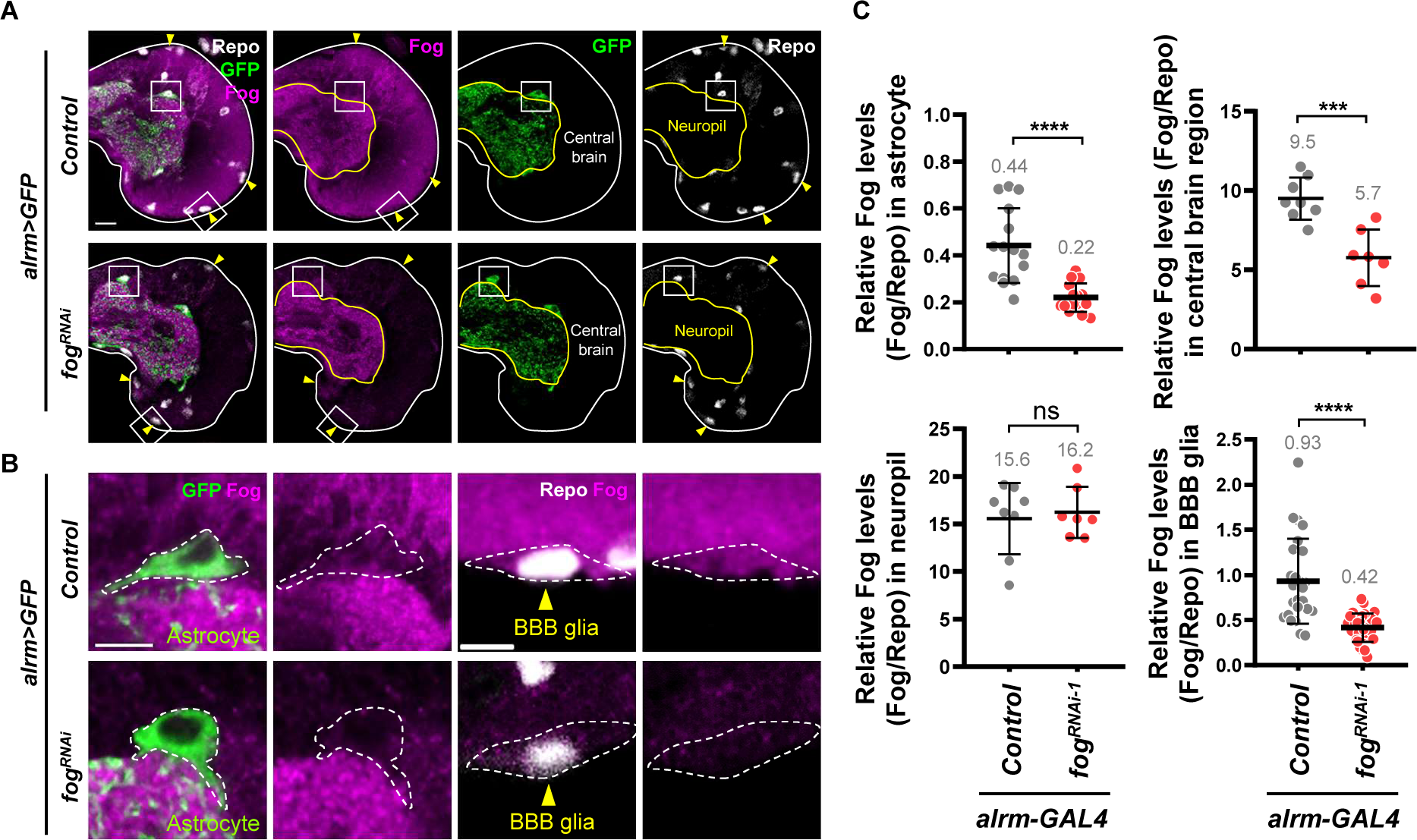
Knocking down *fog* in astrocytes reduces Fog protein in both astrocytes and the larval central brain including BBB glia. **A,** Fog protein levels (magenta) in brain lobes (Repo, white; GFP, green) of different genotypes at 24h ALH in larval brains. White squares mark the region for high magnification of astrocytes and BBB glia in B. White line for the edge of brain lobe; yellow line for the edge of neuropil region. Scale bar, 10 mm. **B,** Fog protein levels (magenta) in astrocytes (Repo, white; GFP, green) and BBB glia (Repo, white, at surface of brain lobes) of different genotypes. Dash outlines, morphology of astrocyte and BBB glia. Scale bar, 5 mm. **C,** Quantification graph of Fog protein levels in astrocytes, central brain regions, neuropil region and BBB glia in control (*alrm>β-gal^RNAi^*) and *alrm>fog^RNAi-1^* groups. Fog protein levels in astrocytes: control, 0.44 ± 0.16, n = 15; *alrm>fog^RNAi-1^*, 0.42 ± 0.06, n = 15. Fog protein levels in central brain region: control, 9.5 ± 1.32, n = 8; *alrm>fog^RNAi-1^*, 5.7 ± 1.78, n = 7. Fog protein levels in neuropil region: control, 15.6 ± 3.76, n = 8; *alrm>fog^RNAi-1^*, 16.2 ± 2.69, n = 7. Fog protein levels in BBB glia: control, 0.93 ± 0.47, n = 25; *alrm>fog^RNAi-1^*, 0.42 ± 0.16, n = 30. P-values were calculated from two-tailed unpaired Student’s t-test for comparison of two samples. ***P < 0.001; ****P < 0.0001.

**Figure S15.**
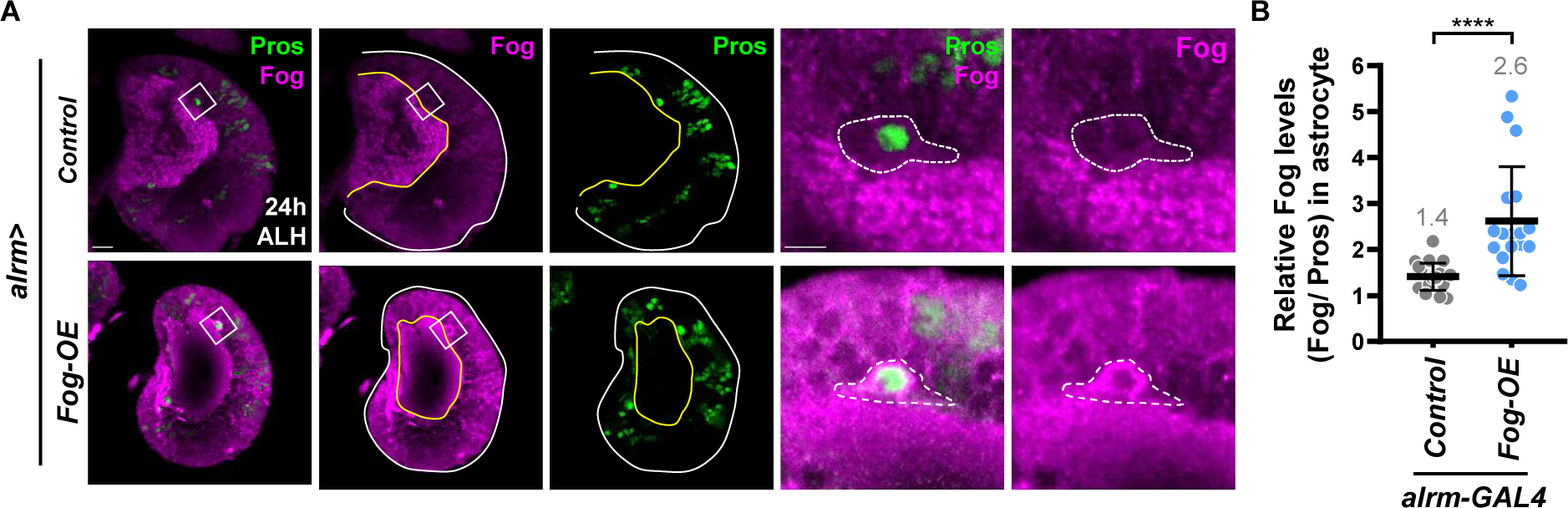
Overexpression of Fog increases Fog protein levels in astrocytes. **A,** Fog protein levels (magenta) in the astrocytes (Pros, green) of different genotypes at 24h ALH in larval brains. White squares mark the region for high magnification. White line for the edge of brain lobe; yellow line for the edge of neuropil region. Dash outlines, astrocyte. Scale bar, 10 mm; 5 mm for images with high magnification. **B,** Quantification graph of Fog protein levels in astrocytes of control (*β-gal^RNAi^*) and Fog-OE conditions. *control* (*β-gal^RNAi^*): 1.41 ± 0.29, n = 21; *Fog-OE*: 2.62 ± 1.18, n = 18. P-values were calculated from two-tailed unpaired Student’s t-test for comparison of two samples. ****P < 0.0001.

**Figure S16.**
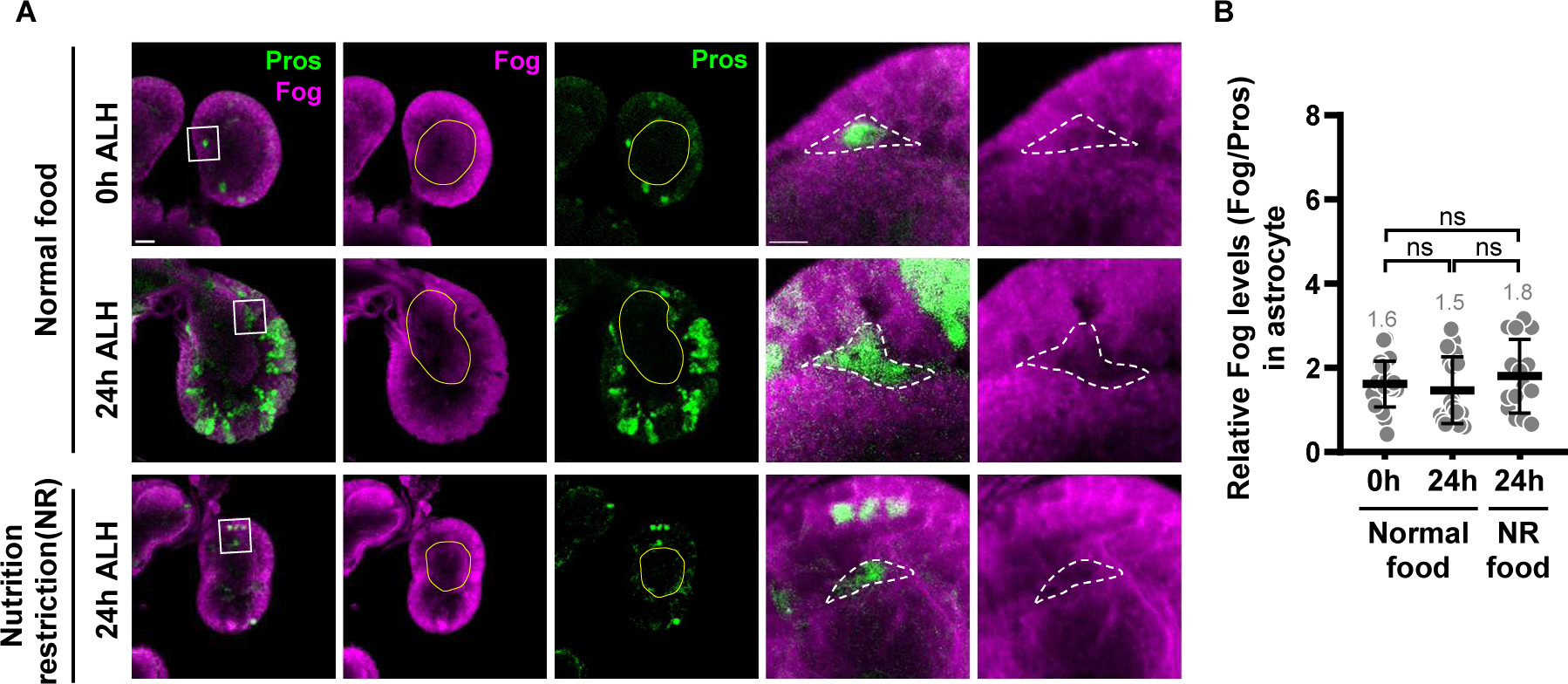
Fog protein expression remains constant during early brain development and is independent of nutrition. **A,** Fog protein levels (magenta) in the astrocytes (Pros, green) of different conditions. White squares mark the region for high magnification. yellow line for the edge of neuropil region. Dash outlines, astrocyte. Scale bar, 10 mm; 5 mm for image with high magnification. **B,** Quantification graph of Fog protein levels in astrocytes in different conditions. Normal food 0h ALH: 1.61 ± 0.55, n = 21; normal food 24h ALH: 1.50 ± 0.79, n = 18; nutrition restriction (NR) food 24h ALH: 1.80 ± 0.87, n = 15. One-Way ANOVA is used for statistics. ns, no significance. The means of analyzed phenotypes were shown above each column.

**Figure S17.**
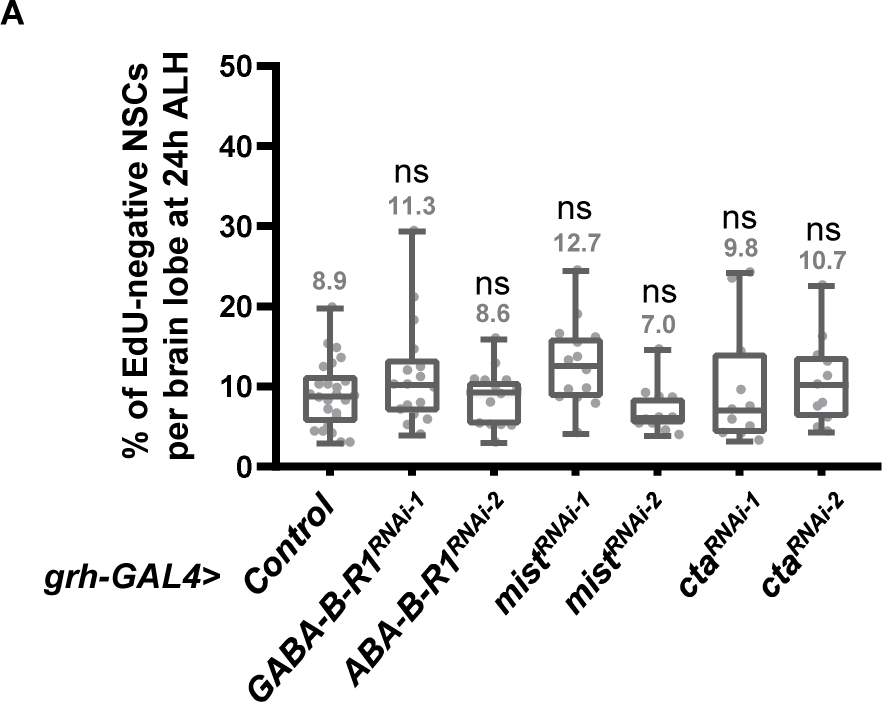
GABA-B-R1 and Mist-cta signaling are not required for NSC reactivation. **A,** Quantification graph of EdU-negative quiescent NSCs in various genotypes : Control (*β-gal^RNAi^*): 8.9 ± 4.2, n = 25; *mist^RNAi-1^*: 12.7 ± 5.3, n = 14; *mist^RNAi-2^*: 7.0 ± 3.0, n = 11; *cta^RNAi-1^*: 9.8 ± 7.6, n = 11; *cta^RNAi-2^*: 10.7 ± 5.5, n = 11. One-Way ANOVA is used for statistics. The means of analyzed phenotypes were shown above each column. ns, no significance. Box-whiskers plots (A, B) show the central lines for median values, edges of box represent upper (75th) and lower (25th) quartiles and whiskers show minimum and maximum values.

**Figure S18.**
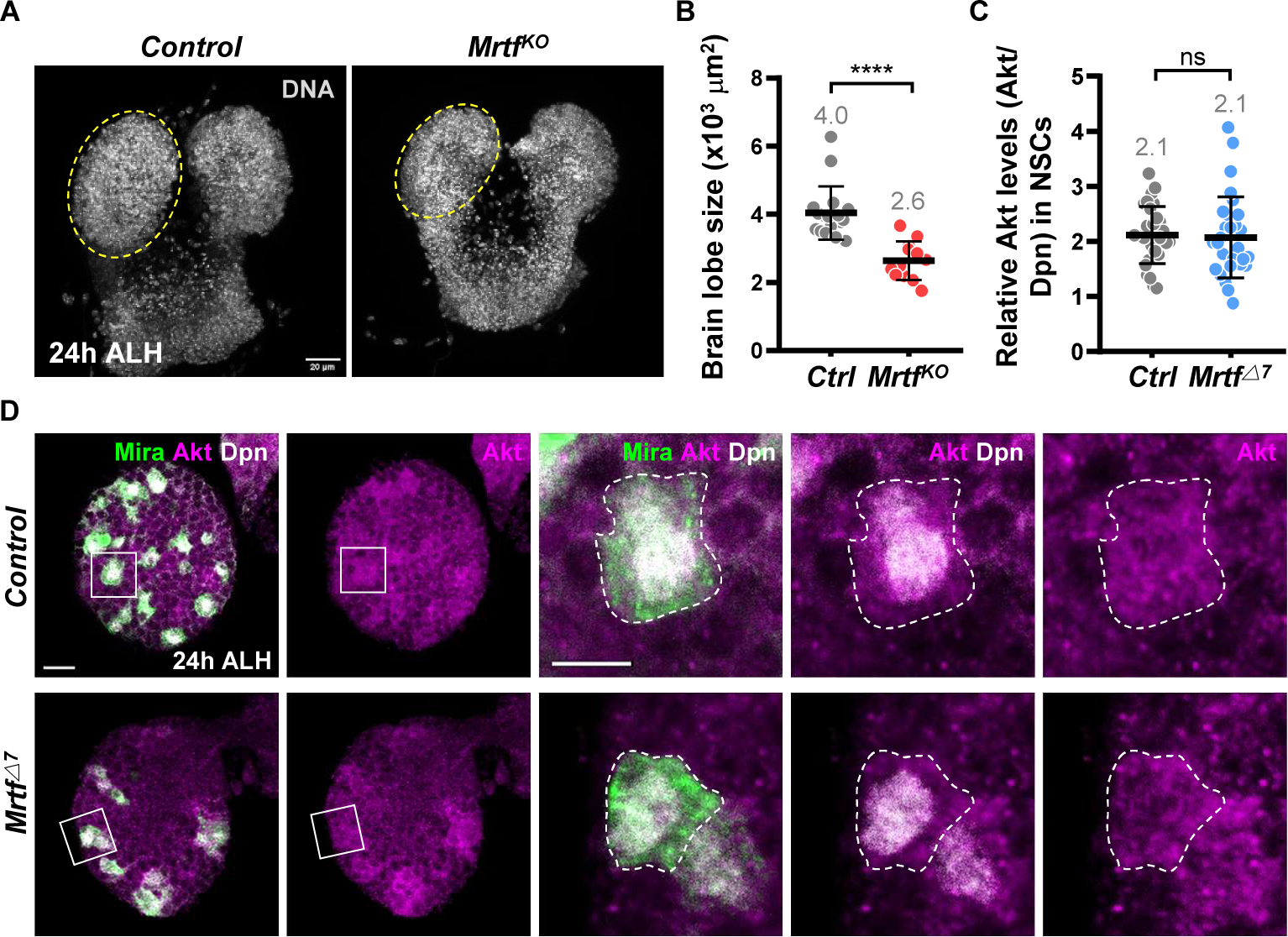
*Mrtf* mutant shows a microcephaly-like phenotype with unaltered Akt levels. **A,** The size of larval brains (DNA, gray) in control (*yw*) and *Mrtf^KO^* mutant at 24h ALH. Dash circles, single brain lobe. Scale bar, 20 mm. **B,** Quantification graph of brain size from Control (*yw*): 4.04 ± 0.78, n = 17; *Mrtf^KO^* : 2.64 ± 0.56, n = 12. **C,** Quantification graph of Akt protein expression in NSCs of control (*yw*) and *Mrtf^KO^* mutants. *control* (*yw*): 2.11 ± 0.52, n = 30; *Mrtf*^△^*^7^*: 2.07 ± 0.74, n = 30. **D,** Akt protein expression (magenta) in the NSCs (Dpn, white; Mira, green) of different genotypes. White squares mark the region for high magnification. Dash outlines, edge of NSCs. Scale bar, 10 mm; 5 mm for image with high magnification.

## Notes

### Competing Interest Statement

The authors have declared no competing interest.

